# Efficient Simulation of Viral Transduction and Propagation for Biomanufacturing

**DOI:** 10.1101/2024.03.30.587435

**Authors:** Francesco Destro, Richard D. Braatz

## Abstract

Viral transduction is a main route for gene transfer to producer cells in biomanufacturing. Designing a transduction-based biomanufacturing process poses significant challenges, due to the complex dynamics of viral infection and virus-host interaction. This article introduces a software toolkit composed of a multiscale model and an efficient numeric technique that can be leveraged for determining genetic and process designs that optimize transduction-based biomanufacturing platforms. Viral transduction and propagation for up to two viruses simultaneously can be simulated through the model, considering viruses in either lytic or lysogenic stage, during batch, perfusion, or continuous operation. The model estimates the distribution of the viral genome(s) copy number in the cell population, which is an indicator of transduction efficiency and viral genome stability. The infection age distribution of the infected cells is also calculated, indicating how many cells are in an infection stage compatible with recombinant product expression and/or with viral amplification. The model can also consider the presence in the system of defective interfering particles, which can severely compromise the productivity of biomanufacturing processes. Model benchmarking and validation are demonstrated for case studies on the baculovirus expression vector system and influenza A propagation in suspension cultures.

**TOC Graphic:** 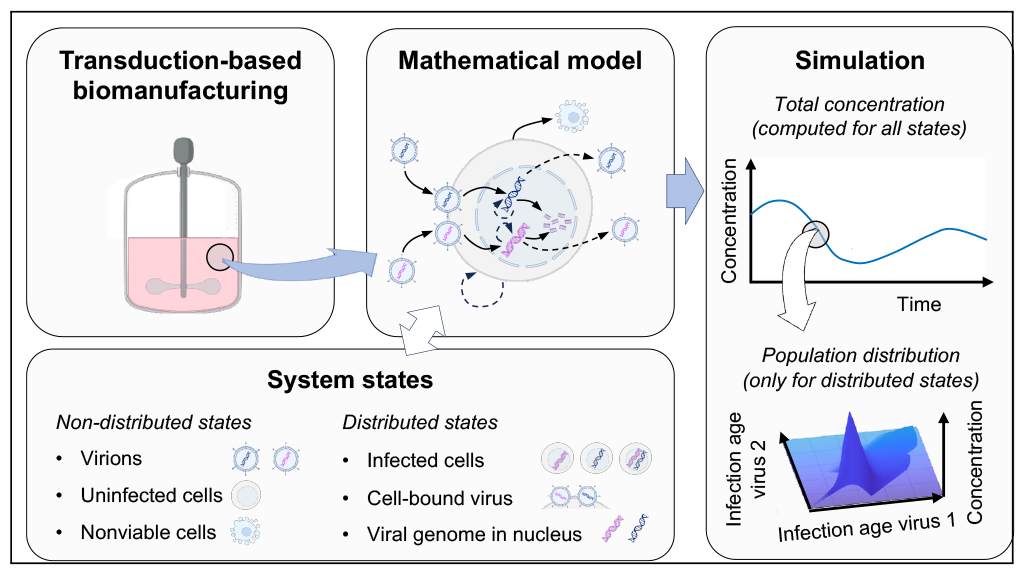

## Introduction

Viral transduction is a primary route for gene transfer to producer cells in biomanufacturing. Several types of viruses are used in academic research and in the industrial practice within transduction-based biomanufacturing platforms. Retroviruses, the most notable example of which are lentiviruses, have been used for producing, among others, virus-like particle (VLP) vaccines^1^ and membrane proteins^2^ in mammalian cells. Transduction-based expression systems based on adenovirus^3^ and herpes simplex virus^4^ have been developed for manufacturing recombinant adeno-associated virus (rAAV), a primary vector for in vivo gene therapy.^5^ The baculovirus expression vector system (BEVS) is an established tool for manufacturing recombinant products with insect cells,^6^ including clinical-grade recombinant vaccines,^7^ VLPs,^8^ and rAAV.^9^ The use of phages as transduction vectors for bacteria has also been explored.^10^ The production of inactivated or live-attenuated virus vaccines for many diseases, including smallpox, polio, measles, and mumps is also based on viral transduction and propagation. ^11^

Since viral transduction consists in the infection of producer cells with recombinant viruses, infection and transduction are here used as interchangeable terms. Scaling up transduction-based biomanufacturing systems to large production poses significant challenges, due to complex viral infection and virus-host dynamics. The optimal multiplicity of infection (MOI) and time of infection (TOI) in batch production depend on a trade-off between preserving high viable cell density and supplying a high level of genetic template for the product of interest. For complex products such as VLPs and viral vectors, it is often necessary to co-transduce multiple viruses, each carrying certain recombinant genes. Optimizing the MOI for each virus requires the evaluation of the most productive combination of recombinant gene copy number within the host. An additional challenge originates in continuous or perfusion processing, which require to consistently maintain target viral titers and viable cell density during viral propagation. Further complications arise from the formation of defective interfering particles (DIPs),^12^ lacking genetic information essential for self-replication and/or for recombinant product manufacturing. In cells coinfected by standard virus (STV) and DIPs, the latter can quickly replicate, interfering with STV propagation. DIPs constitute a serious challenge in biomanufacturing, since they can become predominant over STVs and halt the production of the desired product.

Mathematical simulation is a powerful tool for enhancing the operation of biopharmaceutical processes.^13–15^ So far, few applications of mathematical modeling to enhance transduction-based biomanufacturing systems have been demonstrated, mainly focusing on influenza A.^16,17^Frensing et al.^18^ developed a model for influenza A propagation in continuous bioreactors, using ordinary differential equations (ODEs) to describe STV amplification in the presence of DIPs. The model successfully predicted the insurgence of viral titer oscillations in continuous viral propagation, due to STV/DIP competition (Von Magnus effect ^19^). Since the kinetics of viral infection (e.g., binding, replication, and budding kinetics) strongly vary with the infection age, more advanced models for viral infection need to account for the infection age distribution of the cell population through a system of partial differential equations (PDEs).^12,20^Rudiger et al. ^21^ recently demonstrated that a multiscale PDE model tracking the infection age of infected cells can reproduce experimental data from STV/DIP influenza A systems better than the lumped parameter model developed by Frensing and coauthors. However, the proposed model heavily relies on heuristic parameters fitted to experimental data to describe the intracellular STV/DIP competition. Further, Rudiger et al.^21^ simulated the PDEs of their model by using a finite difference method that causes significant errors in the computation of the infection front due to large numerical diffusion. ^20^ There is a substantial need for (i) a general framework for simulating transduction-based biomanufacturing systems based on different viruses. The underlying mathematical model framework needs to (ii) effectively resolve the coupling between extracellular and intracellular events, (iii) be able to account for the presence and competition among multiple viruses (standard or defective), and (iv) be implemented for batch, continuous, and perfusion systems. Additionally, (v) fast and accurate numeric schemes are needed for this class of PDE models.

This article presents a mathematical simulation framework that addresses these needs. A multiscale mechanistic model and a novel numerical technique are introduced to simulate viral transduction and propagation in batch, continuous, and perfusion biomanufacturing, in the presence of up to two viruses (standard or defective). The framework can simulate any virus/host combination. Three case studies on the BEVS illustrate the use of the model for the design and the optimization of biomanufacturing platforms. The BEVS is chosen as a representative viral system, since the baculovirus infection kinetics are well understood from the literature.^22^ A fourth case study validates the framework with data from the literature on influenza A propagation in presence of DIPs.

## Results and Discussion

### A framework for simulating viral transduction and propagation in biomanufacturing

A novel computational framework for simulating viral transduction and propagation in biomanufacturing systems is presented in Figs. 1–3. The framework is made up of a mechanistic model and of an efficient numerical scheme for solving the model equations. This section summarizes the main features of the model and of the numerics, which are discussed in detail in the Methods section.

**Figure 1.**
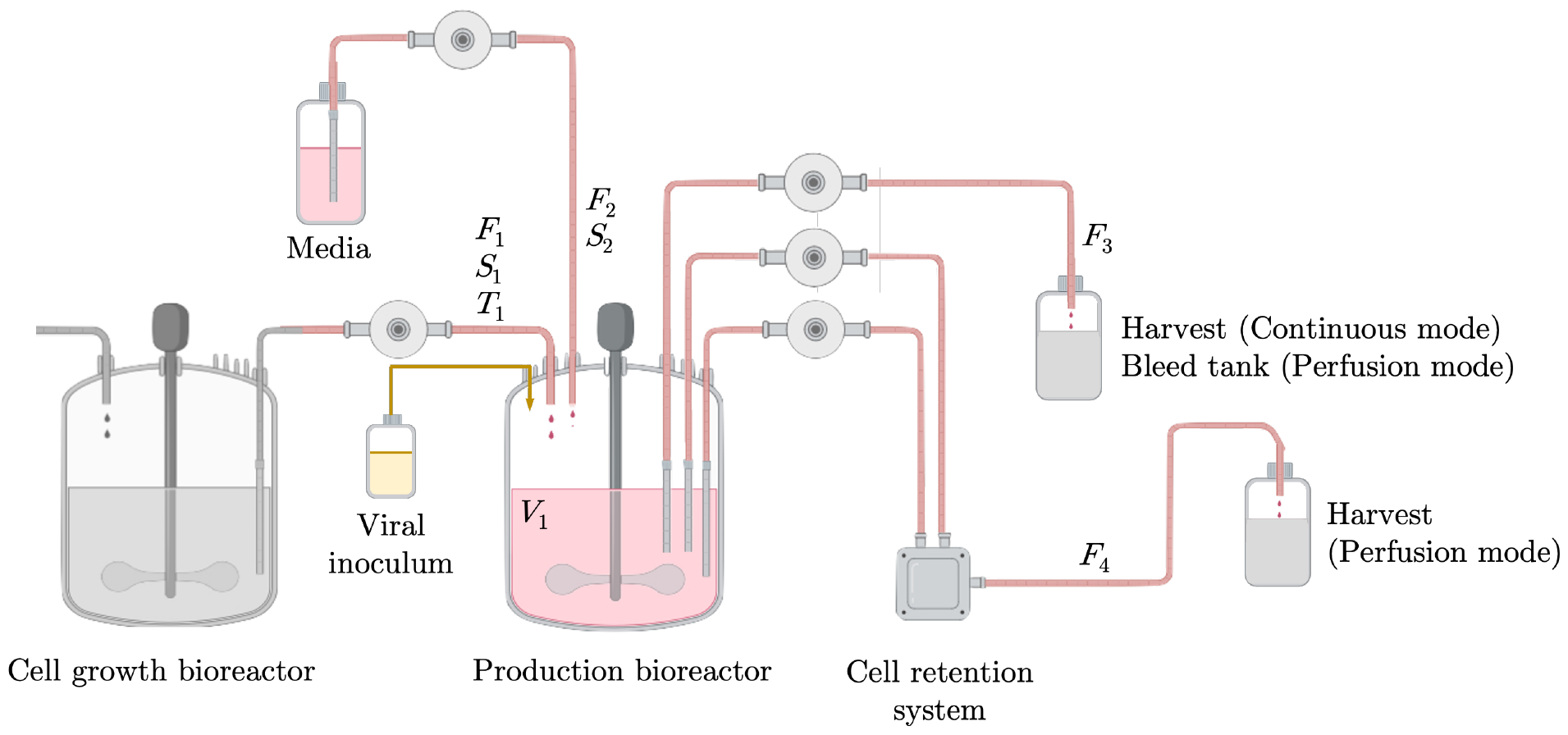
Overview of the biomanufacturing process. The model simulates a production bioreactor in batch, perfusion, or continuous mode; shake-flask experiments can also be simulated. The cell growth bioreactor and the harvest and bleed tanks are not simulated, but are reported here to clarify the input/output structure of the model when run in continuous or perfusion mode.

The model simulates viral transduction and propagation in batch, perfusion, and continuous cell cultures (in bioreactors or shake-flasks; Fig. 1) with up to two viral species, including systems with two STVs or one STV and one DIP. The phenomena considered by the model equations (Fig. 3) are viral binding to host cells, viral genome trafficking to nucleus, degradation, viral replication, progeny production, DIP generation, and the viral coinfection dynamics. The model supports the genetic and process design of transduction-based biomanufacturing systems by estimating critical variables that cannot be readily measured in real time (Table 1). For each virus in the system, the model computes the distributions of infection age and of viral genome copy number in the cell culture (Fig. 2). The distribution of viral genome(s) copy number predicted by the model directly maps to the copy number of template genome(s) for the product of interest. At the same time, the infection age distribution allows to infer how many cells are actively expressing the gene(s) of interest, considering that recombinant product expression occurs only during certain phases of viral infection. Hence, the combined information from the distribution of infection age and the distribution of viral genome copy number allows to estimate the concentration of cells in the system that are producing the recombinant product, and the gene copy number available in each cell for product expression. Further, the model can aid to understand how given process conditions correlate to DIP generation and propagation. To the best of our knowledge, this is the first in silico framework with these capabilities.

**Table 1:** Model: manipulated variables and system states. All system states are expressed as concentrations over the overall system volume. With reference to Fig. 1: 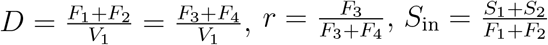 and 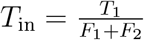 infection age with respect to virus *j* = {1, 2}.

| Symbol | UOM | Description |
| --- | --- | --- |
| **Manipulated variables** |  |  |
| $D$ | h<sup>-1</sup> | Dilution rate: batch: $D = 0$; perfusion/continuous: $D > 0$ |
| $r$ | [-] | Bleeding ratio: perfusion: $0 \leq r < 1$; continuous: $r = 1$ |
| $S_{in}$ | nmol mL<sup>-1</sup> | Substrate concentration in feed |
| $T_{in}$ | cell mL<sup>-1</sup> | Uninfected cell concentration in feed |
| **System states** |  |  |
| $B_{I_j}(t, \tau_j)$ | virus mL<sup>-1</sup> | Virus $j$ attached to $I_j(t, \tau_j)$, for $j = \{1, 2\}$ |
| $B_{C,V_j}(t, \tau_1, \tau_2)$ | virus mL<sup>-1</sup> | Virus $j$ attached to $C(t, \tau_1, \tau_2)$, for $j = \{1, 2\}$ |
| $C(t, \tau_1, \tau_2)$ | cell mL<sup>-1</sup> | Coinfected cells |
| $I_j(t, \tau_j)$ | cell mL<sup>-1</sup> | Cells infected by virus $j$, for $j = \{1, 2\}$ |
| $N_{I_j}(t, \tau_j)$ | vg mL<sup>-1</sup> | Viral genome $j$ in nucleus of $I_j(t, \tau_j)$, for $j = \{1, 2\}$ |
| $N_{C,V_j}(t, \tau_1, \tau_2)$ | vg mL<sup>-1</sup> | Viral genome $j$ in nucleus of $C(t, \tau_1, \tau_2)$, for $j = \{1, 2\}$ |
| $S(t)$ | nmol mL<sup>-1</sup> | Substrate |
| $T(t)$ | cell mL<sup>-1</sup> | Uninfected cells |
| $V_j(t)$ | virus mL<sup>-1</sup> | Extracellular virus $j$, for $j = \{1, 2\}$ |
| $W(t)$ | cell mL<sup>-1</sup> | Nonviable cells |

**Figure 2.**
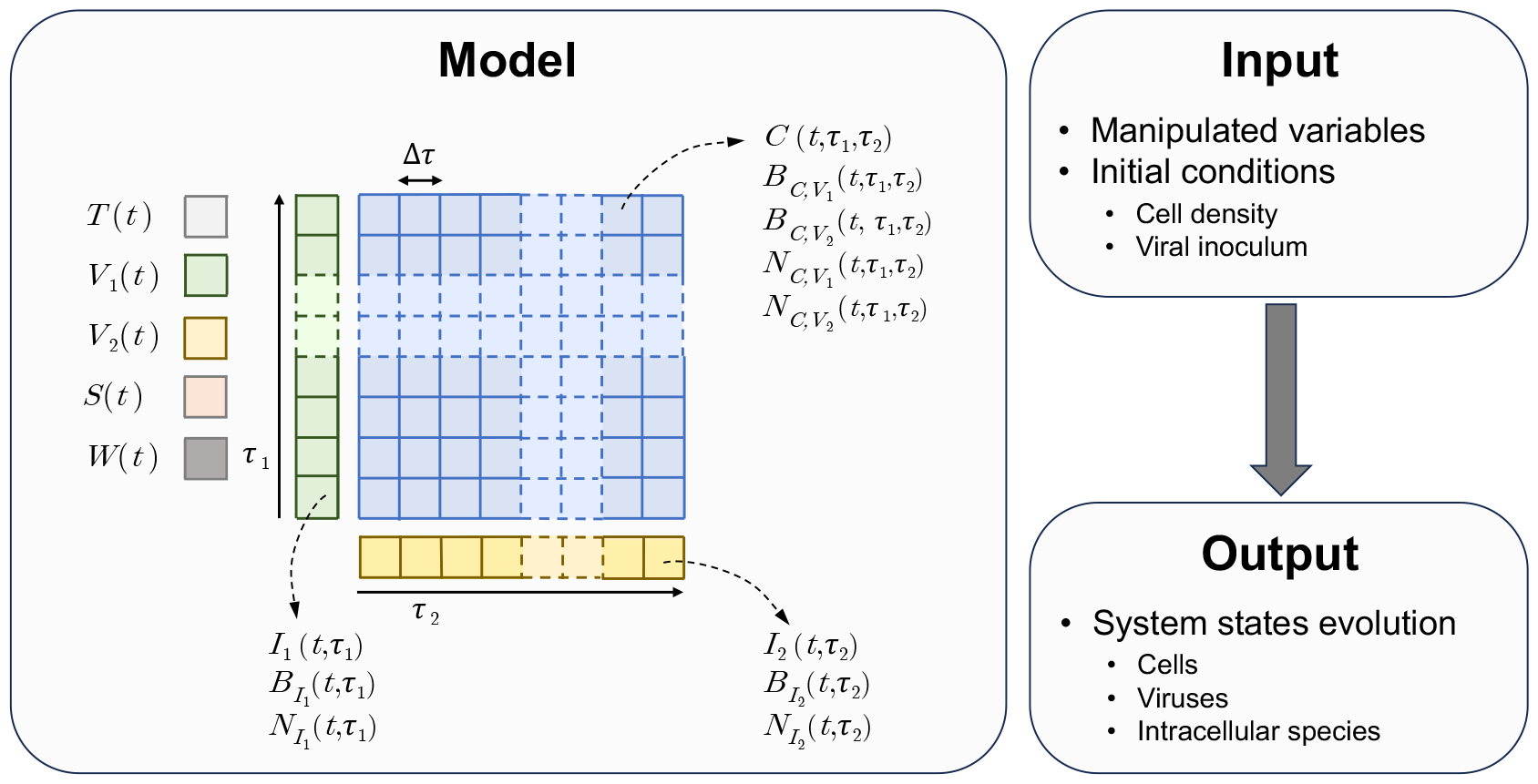
Overview of the model structure. The model inputs are the manipulated variable profiles and the initial value of the system states (Table 1). The model output is the time evolution of the states. The model features 5 ODEs, here represented as standalone squares, and 11 PDEs. Based on the mesh size Δ*τ*, the PDEs are converted into sets of ODEs, here represented as sets of contiguous squares. The concentration of cells infected by only one virus (*I*_1_ and *I*_2_) and the respective amount of surface-bound virus (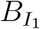 and 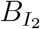) and of nuclear viral genome (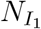 and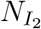) are distributed with respect to one infection age (respectively, *τ*_1_ and *τ*_2_). The concentration of coinfected cells (*C*) and the respective amount of surface-bound virus (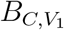 and 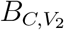) and nuclear viral genome (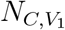and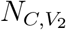) are distributed with respect to both *τ*_1_ and *τ*_2_.

**Figure 3.**
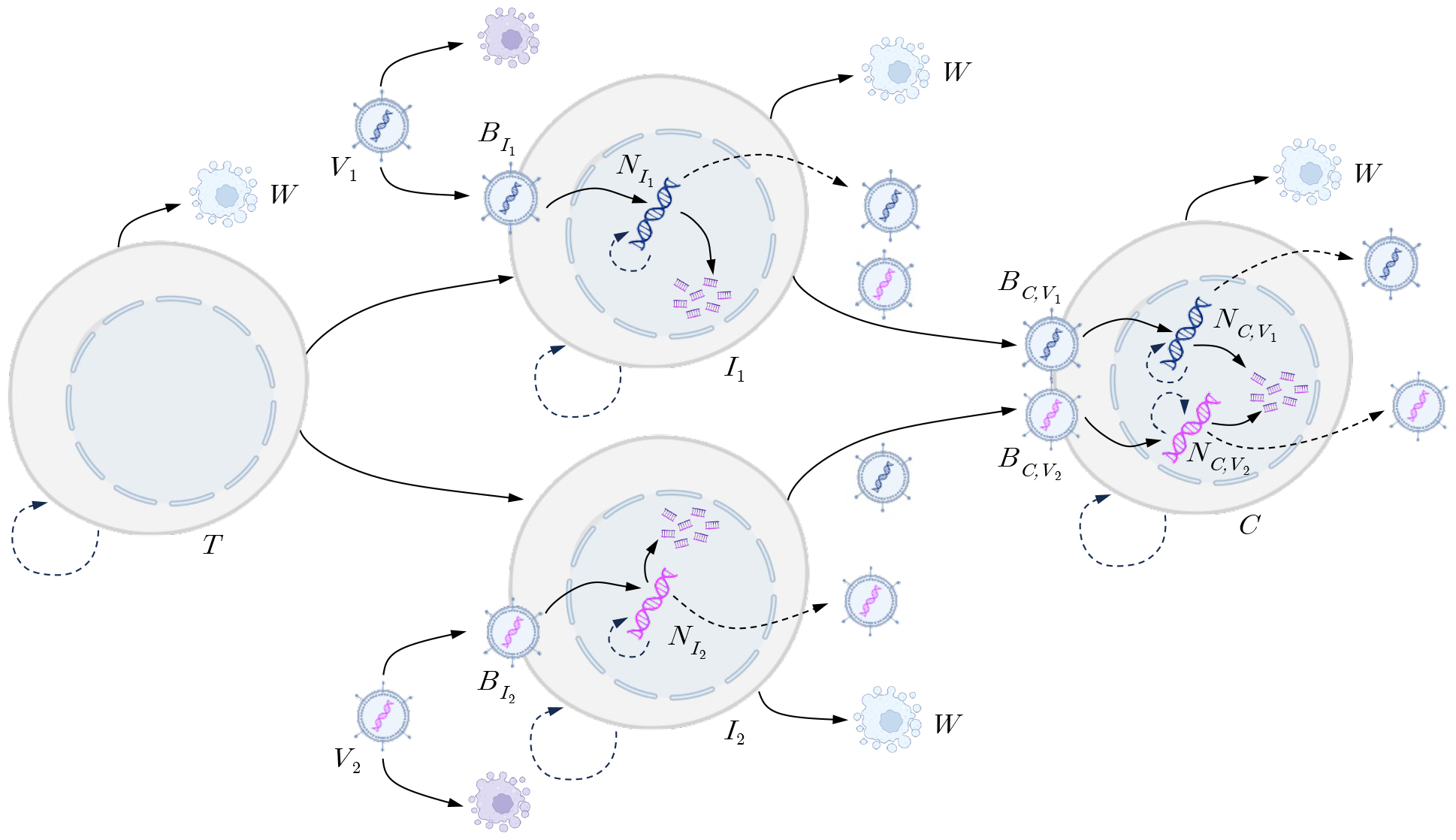
Extracellular and intracellular phenomena considered by the model: cell growth, apoptosis and cell lysis, virus binding, virus transport to nucleus, viral genome replication, viral budding, degradation of extracellular virus and of viral genome. Viral degradation due to rerouting to lysosomes during trafficking to nucleus is also considered by the model, although it is not reported here for conciseness. The legend of the symbols denoting the system states is reported in Table 1.

The model offers a reconciliation between the intracellular and extracellular compartments, which are solved simultaneously with a fast (Table S1) numeric approach that minimizes the detrimental effect of numeric diffusion (Figs. S1–S2). The distributions of infection age and viral genome copy number computed by the model are leveraged for computing the kinetics of the steps of viral infection that are affected by the infection age and by the viral genome copy number, such as viral binding, progeny release, and host viability decay (Fig. 4). In the presence of multiple viruses within a cell, the respective infection ages and genome copy numbers are used to simulate the competition between the two viruses for replicating within the host and for producing progeny (Eqs. S25–S26). This architecture allows the simulation of systems with any host/virus combination. Both RNA and DNA viruses with either single-stranded or double-stranded genomes can be considered, as well as viruses infecting hosts through either endocytosis or membrane fusion. Although the model applications are more interesting for systems that involve active viral replication and propagation, the computational framework also can be leveraged for platforms in which no viral replication occurs (e.g., stable integration with lentivirus). Model limitations that should be considered when using the simulation framework are discussed in the Supporting Information.

**Figure 4.**
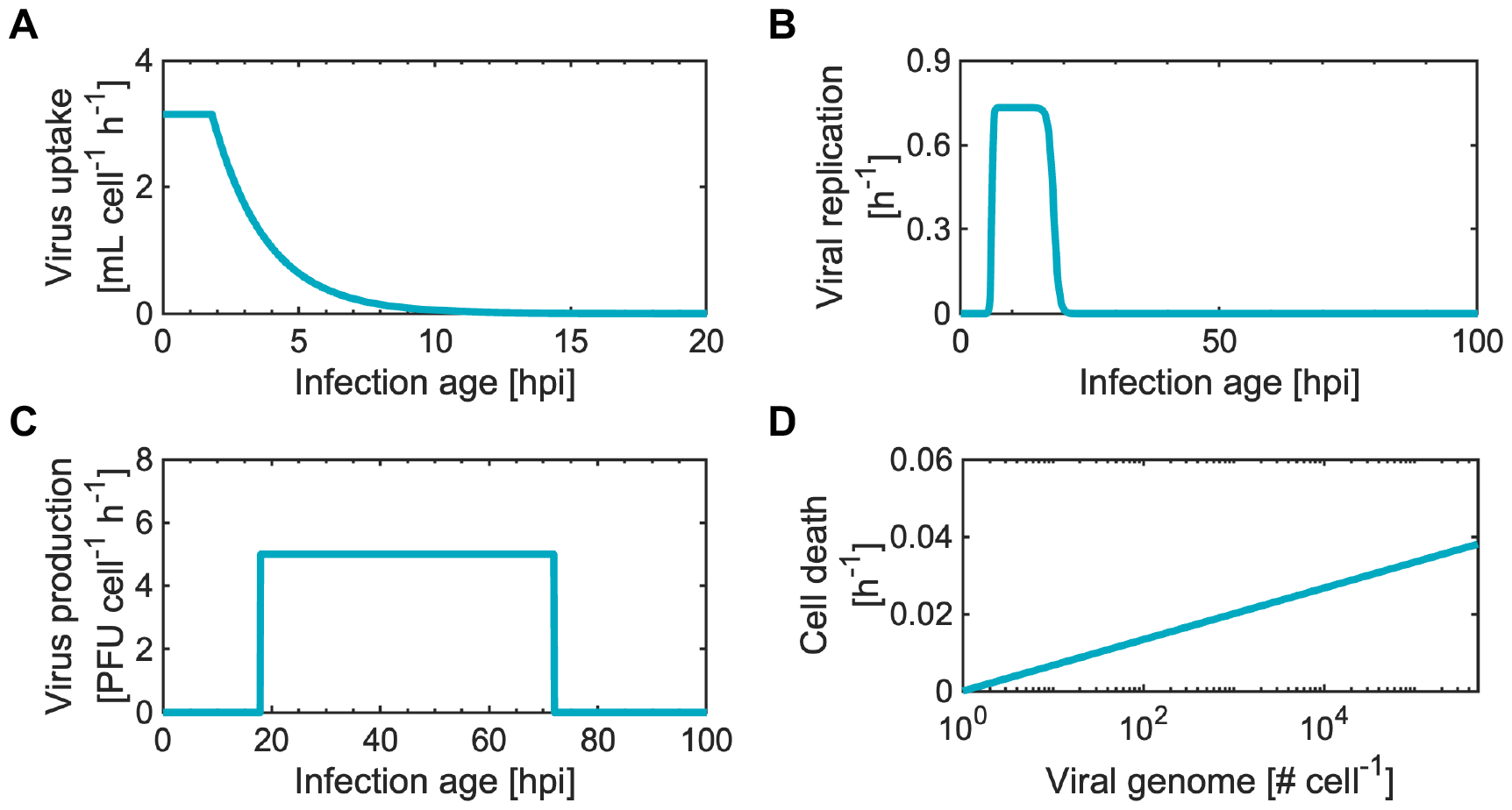
Baculovirus infection kinetics: dependency on cell infection age of (A) viral uptake (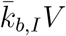; Eq. 3, with viral titer *V* = 5 PFU per cell), (B) viral replication 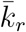; Eq. 11), and (C) viral progeny production 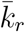; Eq. 6); (D) effect of the viral genome copy number in the cell nucleus on the cell death rate 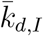; Eq. 4). The full list of parameters for the baculovirus infection kinetics is in Table S3.

The model features 16 equations and 64 parameters to describe the most general scenario of co-transduction from two different recombinant viruses (Table S2). Most model parameters are available in the literature for vectors commonly used in biomanufacturing. The number of parameters reduces to 23 for co-transduction from viruses of the same type that differ only in the recombinant cassette (Table S4, Case study 3), and to 24 to simulate systems with an STV and a DIP of the same type of virus (Table S5, Case study 4). When only one virus is in the system, the model is reduced to 7 equations and 22 parameters (Table S3, Case studies 1–2). The next sections showcase applications of the computational framework in biomanufacturing systems based on the baculovirus expression vector system and on influenza A propagation in presence of DIPs. The proposed numerical scheme is benchmarked against approaches from the state of the art.

### Model Input

#### Case study 1: Scaling up the baculovirus expression vector system

In the BEVS, recombinant baculoviruses deliver the genes for a product of interest to producer cells, usually from the Sf9/Sf21 lines derived from *Spodoptera frugiperda*.^23^ Within the recent interest in increasing the global production capacity for advanced biotherapeutics, scaling up BEVS-based biomanufacturing systems has become a critical objective.^24^ However, the determination of the optimal MOI, cell density, and nutrient concentration at the time of infection in large bioreactors is a challenging task. Due to the high cost necessary for producing the viral inoculum, low-MOI inocula are resorted to at large scale. As a result, the whole cell population in the bioreactor becomes infected only through several viral propagation cycles, up to 3–5 days after viral inoculation. ^25^ Meanwhile, uninfected cells grow and consume the nutrients present in the system. The novel simulation framework presented in this work captures this delicate interplay between MOI, cell growth, and nutrient consumption, allowing to predict the dynamics of baculovirus propagation at very low MOIs (≪ 1) that result in several viral propagation cycles. Figure 5 shows model simulations for batch experiments inoculated at MOI = 0.001, 0.1, and 1 plaque-forming units (PFU) per cell. The presence of only one type of (standard) recombinant baculovirus is considered, assuming that no other standard nor defective viruses are in the system. The simulations are obtained using the kinetic parameters for baculovirus infection estimated in a recent study,^22^ which carried out a comprehensive parameter validation with several experimental datasets. For MOI = 1, the infective virus from the inoculum quickly infects a large portion of the cell population (Figs. 5AB). Only when the cells infected by the viral inoculum start producing progeny virus (i.e., approximately 18 hours post infection, hpi) the remaining uninfected cells are quickly infected. Due to receptor down-regulation induced by baculovirus infection, the rate of viral uptake decreases after infection (Fig. 4A). Accordingly, the PFU titer in the system starts increasing soon after that all cells have been infected (Fig. 5C). Only modest glucose consumption is registered for MOI = 1 (Fig. 5D), since baculovirus infection induces cell cycle arrest, reducing (and, eventually, halting) substrate consumption.^23^ For MOI = 0.1 and 0.01, only a small fraction of the cell population is infected by the viral inoculum. Uninfected cells continue to grow (Fig. 5A) and to consume glucose (Fig. 5D) for several days, before than all cells present in the system become infected by progeny virus (Fig. 5B). For inoculation at MOI = 0.001, all the glucose in the system is depleted, due to the larger extent of cell growth compared to higher-MOI conditions. This result shows that the lower productivity per cell, often registered in BEVS-based biomanufacturing at low MOI,^26^ can be influenced by nutrient depletion. Model simulation can be leveraged to optimize the process conditions in terms of MOI, cell density at the TOI, and nutrient reservoirs that allow to increase productivity. Although only glucose consumption is considered in this case study, the extension to additional nutrient species is straightforward. ^27^

**Figure 5.**
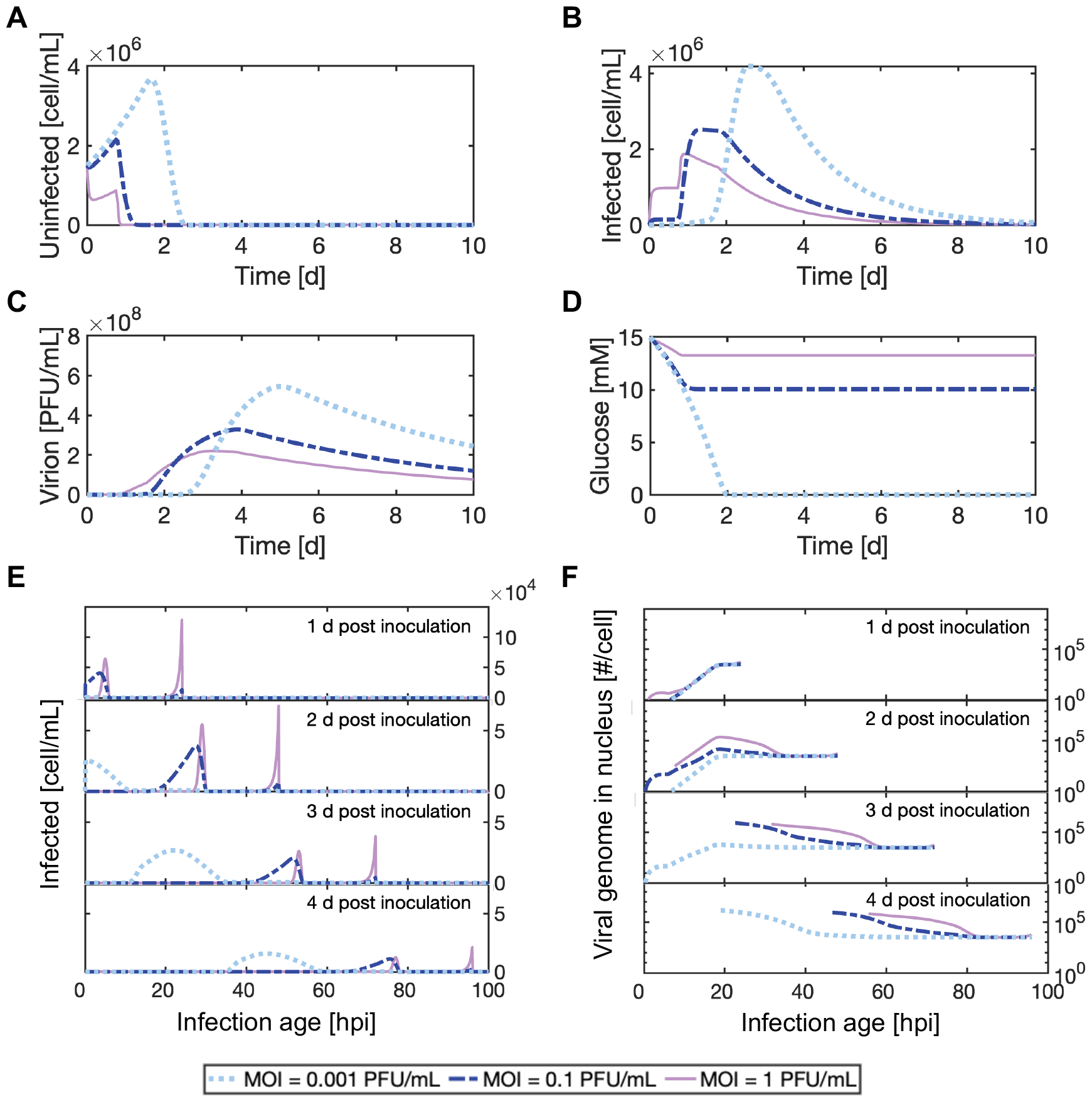
Case study 1: scaling-up the baculovirus expression vector system. Model simulations for three batches inoculated with recombinant baculovirus at, respectively, MOI = 0.001, 0.1, and 1 PFU per cell: (A) uninfected cells concentration, (B) infected cells concentration, (C) virion concentration, (D) glucose concentration, and (E) distribution of infection age and (F) viral genome copy number for the infected cells at 1, 2, 3, and 4 days post inoculation. Plots (E)–(F) are discrete distributions with bin size Δ*τ* = 0.1 hpi. The additional model outputs from Table 1 are not reported for conciseness. Complete simulation settings are in Table S6.

Further, the model provides insights on the infection age of the cells in the system, breaking down the total concentration of infected cells (Fig. 5B) into an infection age distribution (Fig. 5E). One day after viral inoculation, the infection age distribution for MOI = 1 shows a peak corresponding to the cells infected by the inoculum, and a shorter peak associated to the cells infected by the viral progeny. For MOI = 0.01 and MOI = 0.001, the peaks corresponding to the infection wave generated by the viral inoculum are much shorter than for MOI = 1. The cells infected by the progeny virus, instead, present wider distributions of infection age, since more time is needed in these scenarios for infecting the whole cell population (Figs. 5AB). Compared to 1 day after inoculation, the infection age distribution moves forward in the infection age axis at 2, 3, and 4 days after inoculation, while the cell concentration decreases, due to cell death induced by baculovirus infection (Fig. 5E). The corresponding distribution of viral genome copy number in the nucleus of infected cells (Fig. 5F) shows that, at the end of viral replication (≈ 18 hpi), the infected cells achieve approximately a level of viral DNA within the range of 5*×*10^4^–5*×*10^5^ copies per cell, as found in the literature.^28^ An even higher viral DNA copy number is achieved in the cells infected in the late phase of the batch, when the system presents a high titer of progeny virus. This is, for instance, the case for cells with infection age 20 hpi at 4 d post inoculation (Fig. 5F).

However, very few cells achieve these conditions (Fig. 5E), highlighting the importance to consider the infection age and viral genome distributions altogether.

No literature model for the BEVS or general model for viral systems can provide the insights described in this section. Models that do not account for the infection age distribution^18,29^ fail to adequately describe experiments in the BEVS with MOI ≪ 1, since they cannot properly reproduce the viral infection dynamics (Fig. S3). Model benchmarking shows that the novel numerical methodology introduced in this work is successful in accurately tracking the baculovirus infection dynamics (Fig. S1). On the contrary, finite difference approaches, traditionally used for computing infection age distributions in viral systems models from the literature,^12,21^ fail to accurately track the infection front, due to strong numeric diffusion (Fig. S1).

#### Case study 2: Continuous processing in transduction-based biomanufacturing platforms

Continuous and perfusion processing are acquiring importance as innovative approaches for scaling up transduction-based biomanufacturing platforms.^30,31^ Perfusion provides producer cells with fresh media, counteracting the nutrient depletion issues that can be encountered in batch operation. Continuous processing goes even further, by exploiting viral propagation to infect a stream of producer cells that continuously enter a production bioreactor (Fig. 1). Successful establishment of fully-continuous operation can theoretically lead to a system that only requires a fresh media stream to continue manufacturing large quantities of high-value recombinant products. Specific challenges originate in designing the operation of continuous and perfusion processing, which require to consistently maintain target viral titers, viable cell density (VCD), nutrient levels, and high product yield during viral propagation. Figure 6 shows model simulation for 10 days of continuous operation in the BEVS, to compare scale-up strategies based on continuous processing with batch scale-up (Fig. 5). Also in this case, only one type of recombinant baculovirus is present in the system, neglecting the presence of other standard or defective viruses. Simulations are carried out for viral inoculation at MOI = 0.001, 0.1, and 1 PFU per cell. The profiles of uninfected (Fig. 6A) and infected (Fig. 6B) cells density, of PFU titer (Fig. 6C), and of glucose concentration (Fig. 6D) in batch and continuous operation are very similar for the first 3 days after inoculation. Afterwards, different dynamics arise. Interestingly, glucose depletion is achieved about 2 days after inoculation with MOI = 0.001 also in continuous mode (Fig. 6D), demonstrating that a quantitative analysis should be conducted to guarantee that sufficient nutrients are available to the cells, even for perfusion and continuous operation. A steady state is achieved about one week after inoculation. For all the considered MOIs at inoculation, the same steady state is reached, including for the distribution of infection age (Fig. 6E) and of nuclear viral genome (Fig. 6F) for the infected cells. The model predictions provide crucial information to select manipulated variables profiles (Table 1) and to guide baculovirus genetic designs to optimize the process yield. Notably, the promoters typically used to express recombinant products in the BEVS are active only during certain infection age intervals. ^32^For instance, the strong p10 and polh promoters, used in several biomanufacturing processes, are active, approximately, only 15–20 to 35–50 hpi.^9,22,32,33^ Hence, the model can be used for developing process and vector designs that maximize the concentration of cells at infection ages compatible with product expression (Fig. 6E), and the copy number of recombinant genome in those cells (Fig. 6F). As for batch operation at low MOI, literature models that do not account for the infection age distribution^18^ fail to adequately simulate continuous manufacturing in transduction-based platforms (Fig. S4), while simulating the infection age distribution through finite difference approaches leads to low accuracy (Fig. S2).

**Figure 6.**
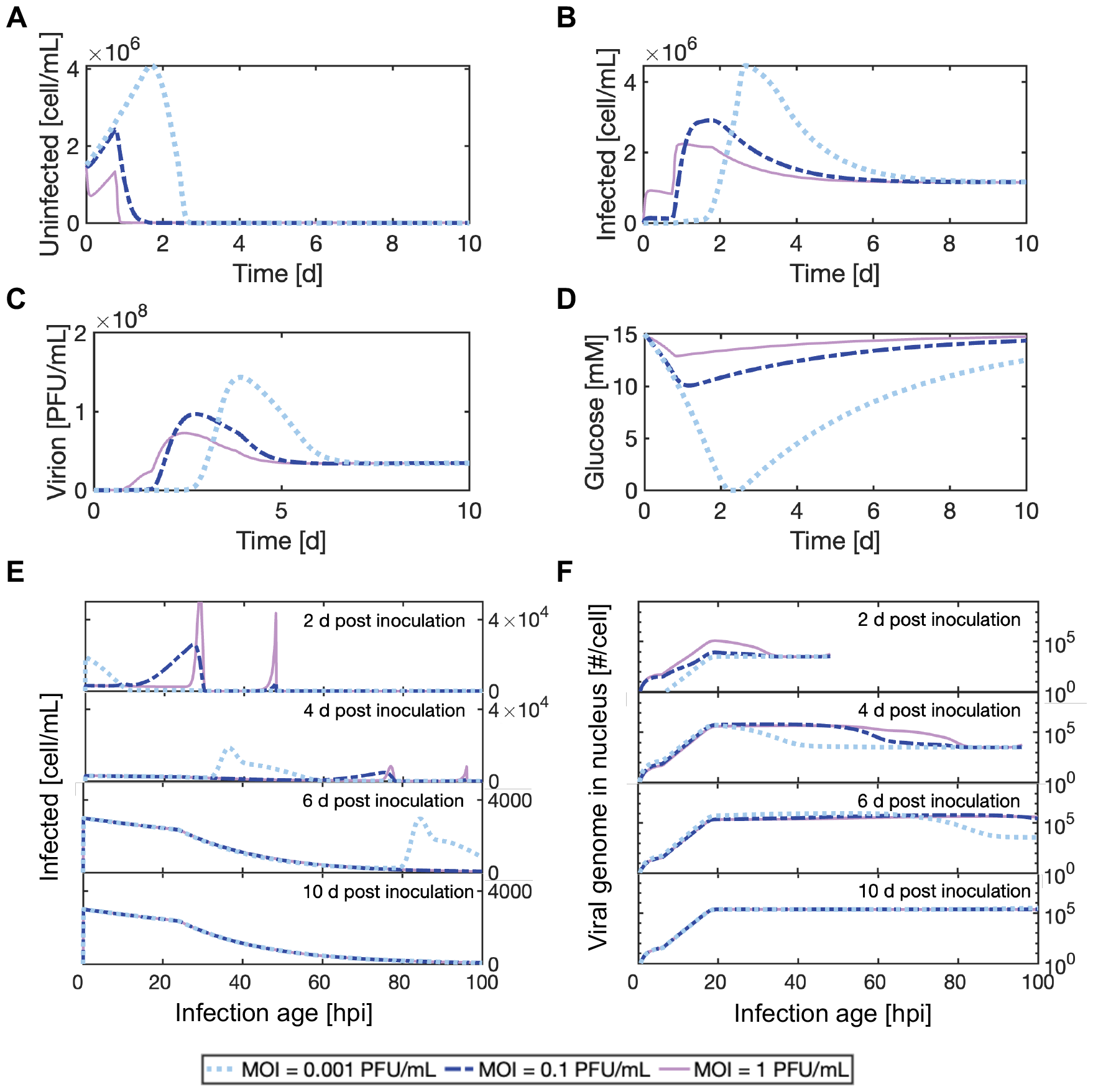
Case study 2: continuous processing in transduction-based biomanufacturing platforms. Model simulations for three runs of continuous operation with the BEVS, in a bioreactor inoculated with recombinant baculovirus at, respectively, MOI = 0.001, 0.1, and 1 PFU per cell: (A) uninfected cells concentration, (B) infected cells concentration, (C) virion concentration, (D) glucose concentration, and distributions of (E) infection age and (F) viral genome copy number for the infected cells at 2, 4, 6, and 10 days post inoculation. Plots (E)–(F) report discrete distributions with bin size Δ*τ* = 0.1 hpi. The additional model outputs from Table 1 are not reported for conciseness. Complete simulation settings are in Table S6.

#### Case study 3: Simulation of viral co-transduction

The genes necessary for recombinant product manufacturing are oftentimes transferred to producer cells through separate viruses.^4,9,33^ The model presented in this work can be used for simulating viral transduction and propagation in presence of two recombinant viruses, to investigate the impact on productivity of different process designs and of genetic modifications to vector and host. Figures 7–8 show a model simulation for a continuous biomanufacturing process based on co-transduction of producer cells by two recombinant viruses (we here refer to co-transduced and coinfected cells interchangeably). Rather than directly using suspended virions, viral inoculation is carried out by introducing infected cells into the bioreactor, as often done in biomanufacturing.^25^ The two viruses, denoted as virus 1 and virus 2, are two baculoviruses that differ only in the recombinant cassette carried within the same genomic backbone. Cells infected by virus 1 and virus 2 are inoculated at, respectively, a 1:100 and a 0.5:100 ratio with respect to uninfected cells. The inoculated cells have an infection age of 40 hpi, hence they start to release progeny virus soon after inoculation. For the first day after inoculation, the uninfected cells are slowly infected by the progeny of the inoculated infected cells (Figs. 7A–D). Most cells infected during this first phase of the process are infected only by either virus 1 or virus 2 (Fig. 7B). At 1 day after inoculation, the infection age distribution of the cells infected by only one virus displays a peak at 64 hpi associated to the viral inoculum, and a distributed wave of more recently infected cells (Fig. 7E). About 1 day after inoculation, the first wave of infected cells starts to produce progeny virus, which quickly infects the whole cell population. Hence, most cells infected later than 1 day after inoculation are coinfected by both virus 1 and virus 2 (Figs. 7B, 7E, 8A–B). Ten days after inoculation, the coinfected cells (Fig. 8C) present the steady-state distribution of infection age expected for baculovirus infection, with a significant decay of viability starting around 24 hpi (Table S4). Interestingly, coinfected cells at 10 days after inoculation show a very small difference of infection age between virus 1 and virus 2, indicating that they are coinfected simultaneously, soon after that they enter the bioreactor, due to the high viral titer (Fig. 7C). The corresponding genome distribution for virus 1 in coinfected cells at 10 days after inoculation is reported in Fig. 8B. The intracellular viral genome concentration reaches the highest level around 18 hpi, as found in the case study with only one virus (Fig. 5E). Cells in infection stages typically associated with recombinant product expression (infection age greater than 15–20 hpi) ^32^ have approximately 50% more viral DNA copies than the average of the entire population of coinfected cells (Fig. 7D). Interestingly, virus 1 (50% more abundant than virus 2 at inoculation) remains predominant in the system after that a steady-state is reached, both as a virion (Fig. 7C) and as viral genome copy number in co-transduced cells (Fig. 7D).

**Figure 7.**
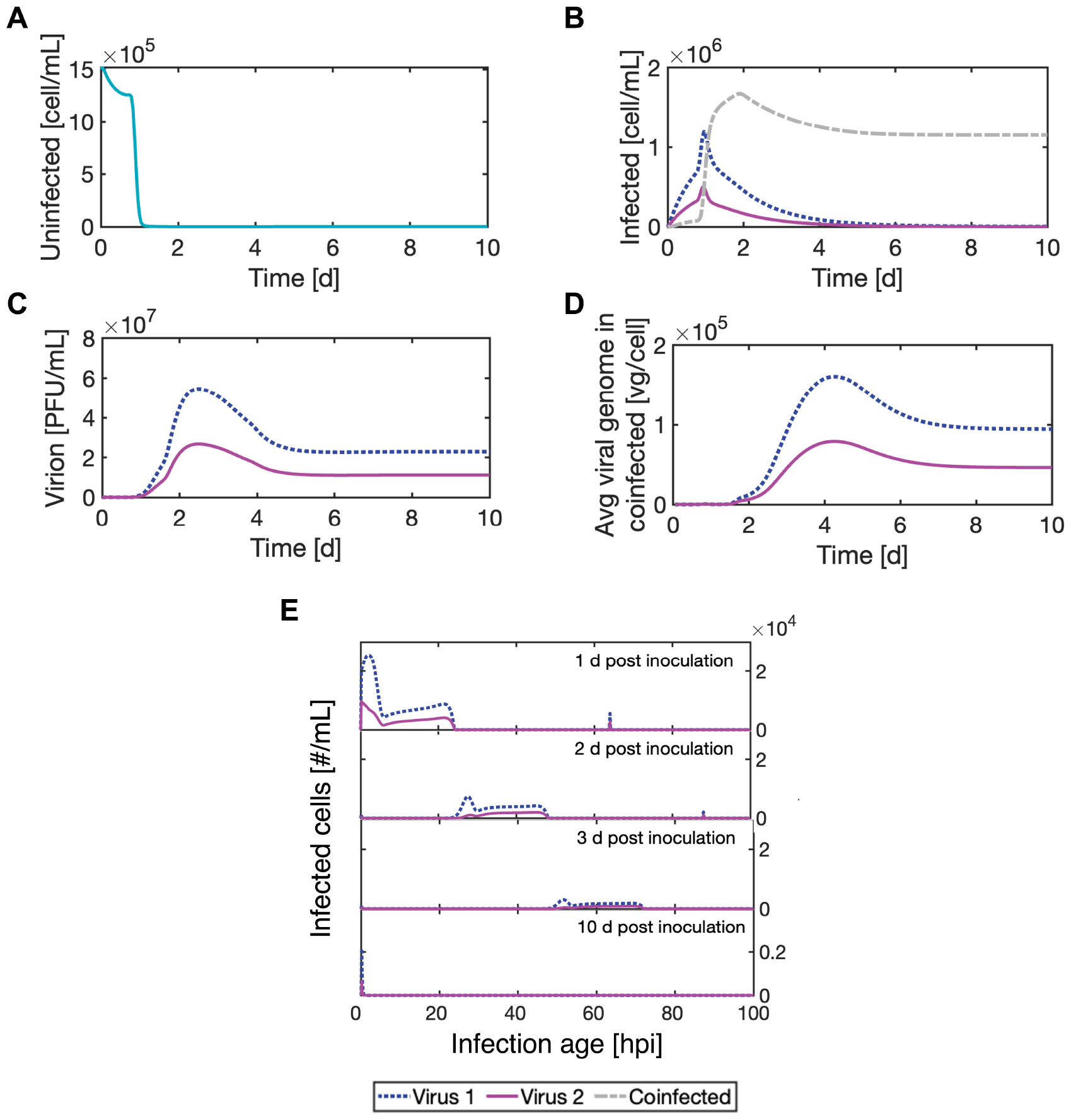
Case study 3: simulation of viral co-transduction in the BEVS (continuous processing). Concentrations of (A) uninfected cells, (B) infected cells, (C) virions, and (D) viral genome in coinfected cells (average); (E) infection age distribution (bin size Δ*τ* = 0.2 hpi) of cells infected by only one virus at 2, 4, 6, and 10 d post inoculation. The complete model output from Table 1 is not reported for conciseness. Complete simulation settings are in Table S6.

**Figure 8.**
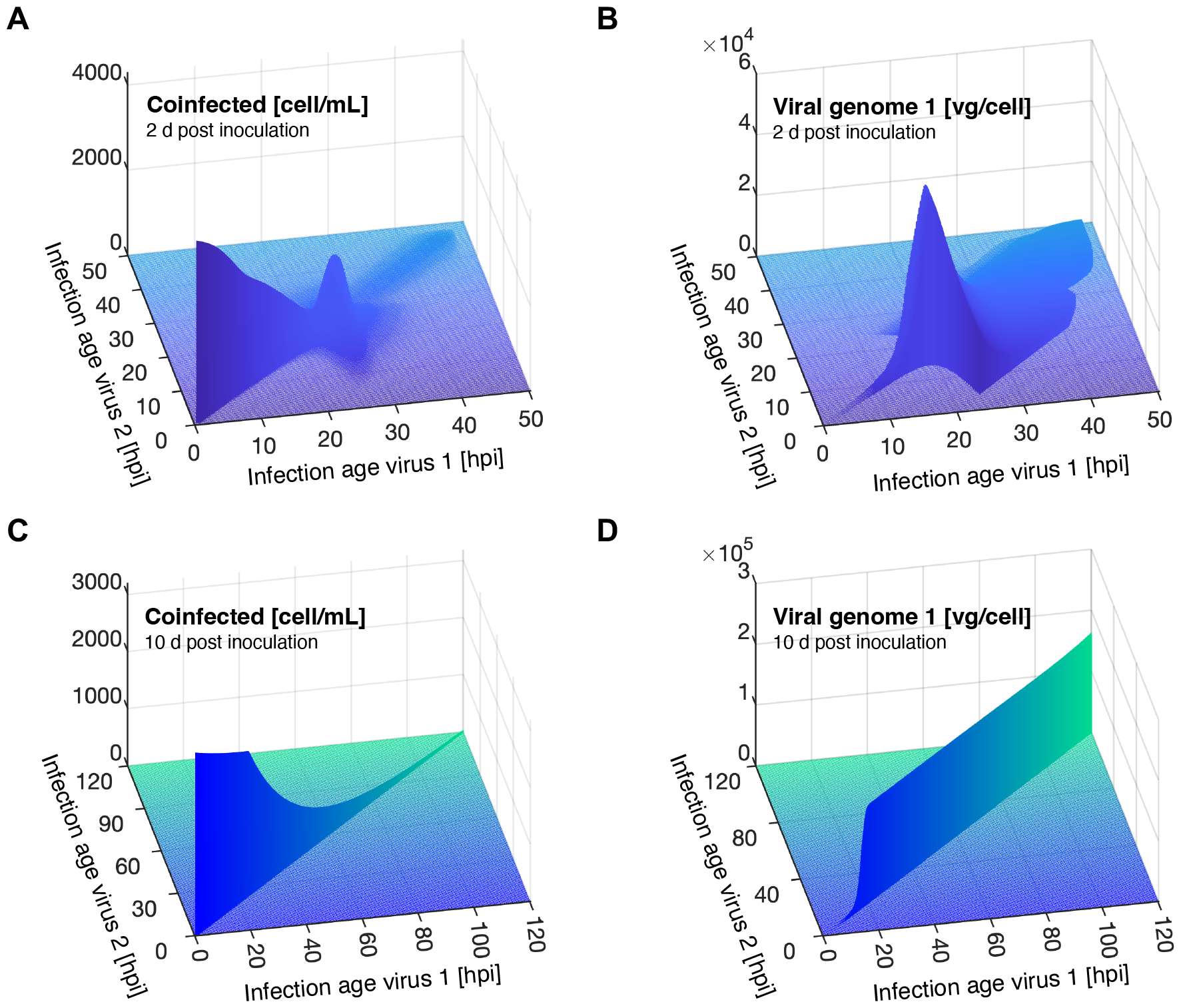
Case study 3: sample distributions of infection age and viral genome for coinfected cells. Two days post inoculation: distributions of (A) infection age and (B) virus 1 copy number. Ten days post inoculation: distributions of (C) infection age and (D) virus 1 copy number. All figures report discrete distributions with bin size Δ*τ* = 0.2 hpi. The viral copy number is not reported for bins with cell concentration lower than 0.1 cell mL^−1^.

All simulations discussed in this and in the previous sections neglect the formation of DIPs and the loss of recombinant cassettes in the baculovirus genome. These phenomena become significant for the BEVS only for higher passage numbers than those considered so far (15–20 passages in batch operation, ^34^ or approximately 10–15 days after viral inoculation for continuous operation^35^). For longer process durations, DIPs and genome loss can destabilize the steady-state conditions that have been shown for continuous processing in the BEVS; further, DIPs can completely hinder the establishment of a steady-state for other viral platforms.^18^The next section discusses simulation of STV/DIP competition through the model presented in this work.

#### Case study 4: Influenza A propagation in presence of defective interfering particles

Viral propagation of most DNA and RNA viruses leads to the formation of DIPs, namely virions lacking a large portion of viral genome and thus incapable of self replication.^19,36^DIPs can replicate at a faster rate than STVs in STV/DIP coinfected cells, probably due to their shorter genome. ^37^ As a result, STV/DIP coinfected cells produce a larger DIP than STV progeny, and DIPs can quickly become predominant during serial passage or continuous processing within viral systems. Influenza is one of the viruses for which DIPs have been characterized most extensively.^18,21,38,39^ DIPs have been identified as promising candidates for influenza antiviral therapy.^40^ However, the formation of DIPs is an adverse event during the production of live attenuated influenza vaccines in immortalized cells cultures, since DIPs cause severe titer oscillations that are detrimental to productivity.^18^ Rudiger et al. ^21^ recently reported experimental data for influenza A propagation in suspension-adapted Madin-Darby canine kidney (MDCK) cells in presence of DIPs. In batch experiments, MDCK cells were inoculated with different combinations of MOIs of influenza A (H1N1) STVs and DIPs DI244,^41^ containing a deletion of the gene coding for the polymerase basic protein 2 (PB2). Twelve experiments were carried out with a full factorial design, using three levels for the STV MOI (0.001, 3, and 30 PFU/mL), and four levels for the DIP MOI (0, 0.001, 3, and 30 PFU/mL). Experimental measurements were collected for viral titers of STVs and DIPs, VCD, fractions of infected and apoptotic cells, and intracellular viral RNA copy number for STV and DIP. The data reported by Rudiger et al. ^21^ are used for validation of the model introduced in this work. As detailed in the Methods section, the model implementation for the considered influenza A STV/DIP system has 19 equations and 24 adjustable parameters (Table S5). While 13 parameters are directly retrieved from the literature, the remaining 11 parameters are estimated from experimental measurements of infectious STV and total DIP titer (Fig. 9), VCD (Fig. S6), and intracellular viral genome copy number (Fig. S5). Only 8 of the 12 experiments reported by Rudiger et al. ^21^ are used for parameter estimation; the remaining 4 experiments are retained as validation dataset. An excellent estimation is achieved for the STV and DIP titers for all the experiments (Fig. 9 and Table S7). The STV and DIP titers are the key variables for this process, since they represent the products of interest for the manufacturing of, respectively, influenza vaccines and DIP antiviral therapy. The model is successful in predicting also the VCD (Fig. S6) and the intracellular level of standard and defective viral genome (Fig. S5). For the latter, the measurement error can be very significant, since the intracellular level of viral genome is inferred through qPCR and VCD signals with standard deviation of, respectively, 55% and 40%. ^42^ Measurements of infected cells concentration are further used for validation (Fig. S7). The worst fit for all measurements is achieved in the experiments at STV MOI equal to 0.001, for which a small measurement error in the inoculated MOI has the largest effect on the experimental dynamics. Further, constant offsets in viral titers and intracellular viral genome measurements, more noticeable at low titer, are expected due to background noise.^21^Hence, the model is successful in predicting the impact of DIPs onto STV propagation in all the experiments, both at the extracellular and at the intracellular level.

**Figure 9.**
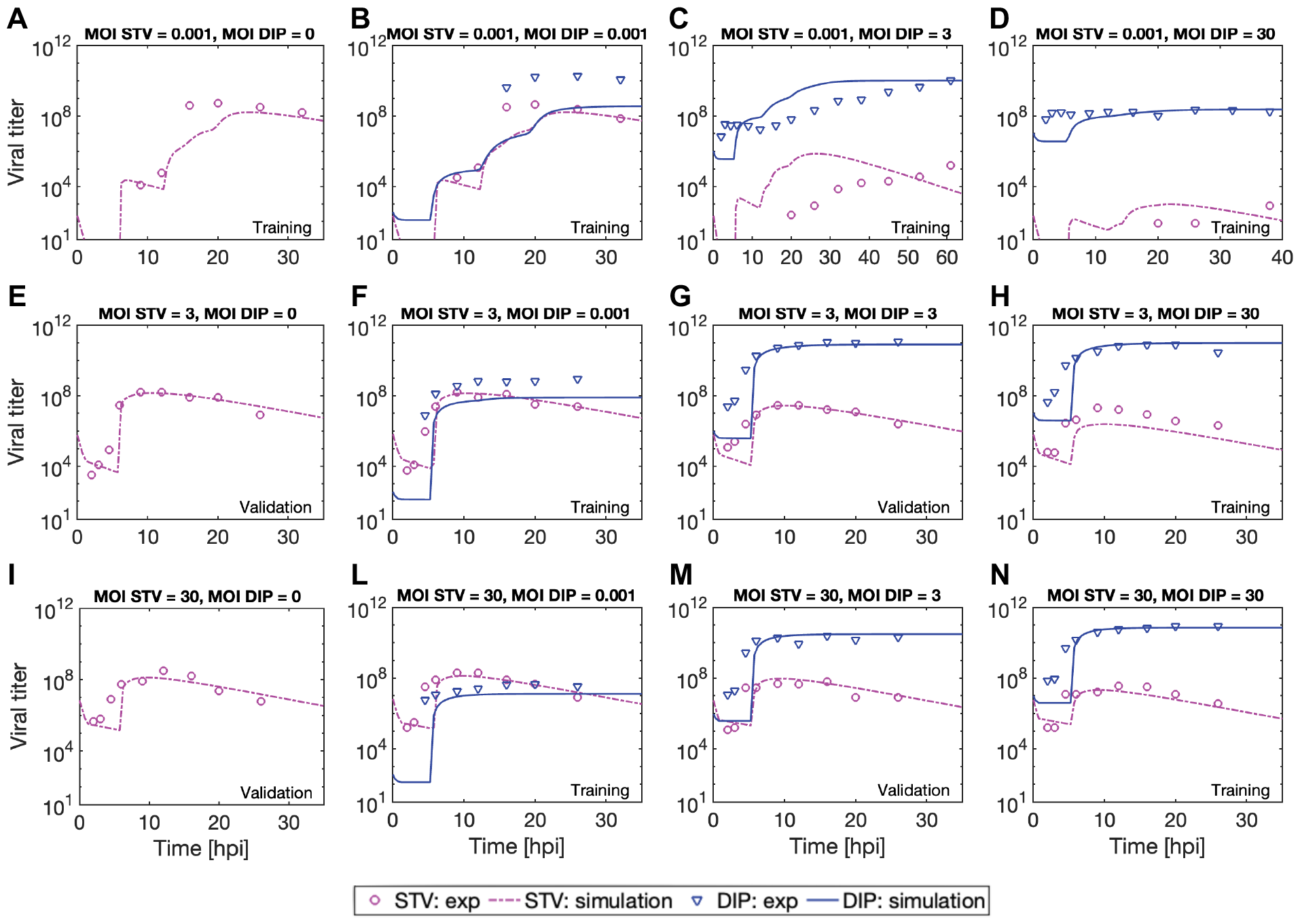
Case study 4: influenza A propagation in presence of defective interfering particles. Model predictions vs. experimental measurements for the titers of influenza A STV (infectivity assay) and DIP (real-time quantitative PCR) in 12 batches. Each batch is inoculated at the reported STV MOI, measured through infectivity assay, and DIP MOI, measured through infectivity assays on MDCK cells genetically modified to express PB2 (“active DIP titer” approach^46^). Data from batches (A)–(D), (F), (H)–(J), and (L) are used for parameter estimation, and (E), (F), (I), and (K) are used only for model validation. Experimental data are from Rudiger et al. ^21^

The most advanced model in the state of the art of influenza A STV/DIP systems (and, to the best of our knowledge, of viral transduction and propagation systems) has been presented by Rudiger et al., ^21^ in the same article that reports the experiments here considered.^21^The model by Rudiger et al.^21^ contains 132 differential equations and 73 parameters, which summarize and improve a set of models proposed in recent years.^18,42–45^ Table S7 compares the residual sum of squared errors (RSS) and the Akaike information criterion (AIC) for the model presented in this work with those for the model by Rudiger et al. ^21^ RSS and AIC are calculated on experimental measurements of the key performance indicators for the process: viral titer of (i) STV and (ii) DIP, intracellular viral genome level of (iii) STV and (iv) DIP, and (v) VCD. Rudiger et al. ^21^ discussed model validation also for experimental measurements of three mRNA transcripts that intervene during influenza propagation. mRNA measurements are not included in the RSS and AIC computation here, since mRNA and other non-crucial intermediates are lumped together in our model, to reduce the computational burden. The model introduced in this work achieves a better fit than the model by Rudiger et al.^21^ in terms of both RSS and AIC. This result is even more significant considering that the Rudiger model has 113 differential equations and 49 parameters more than our proposed model, and that Rudiger et al. ^21^ estimated 8 more parameters on the same experimental dataset, and used all 12 experiments for parameter estimation. The Supporting Information provides a detailed discussion of the distinctive features of our model that allow this groundbreaking performance. In essence, the computation of the two-dimensional infection age and viral genome copy number distributions (Fig. 2), as introduced in this work, allows to capture the intracellular STV and DIP competition at an unprecedented level of detail. The novel numerical approach is an enabling feature of the proposed simulation framework, since it allows the propagation of several distributed states in a low computational time (Table S1).

## Methods

### Mathematical Modeling

The model developed in this work describes viral transduction and propagation for up to two viral species in a well-mixed system, operated in batch, continuous, or perfusion mode (Figs. 1–3). The model is based on mass balances over the system volume for the species listed in Table 1. The system volume is considered constant in all equations. In this section, all distributed states are introduced as continuous distributions, while the main text reports all distributions in discrete form. Continuous distributions are represented using lowercase letters for symbols, while discrete distributions are denoted by the corresponding uppercase letters of the same symbol, with unchanged subscripts and superscripts. Non-distributed states are always reported as uppercase letters. The Numerics section details how continuous distributions are converted into discrete distributions. The remainder of this section describes the model for systems presenting only one viral species (used in Case studies 1–2). The complete model for systems with two viral species is given in the Supporting Information. The models give completely equivalent results when only one virus is present in the system, although the model for systems with only one virus has a much lower computational demand.

The balance for the uninfected cells concentration *T* (*t*) [cell/mL] is

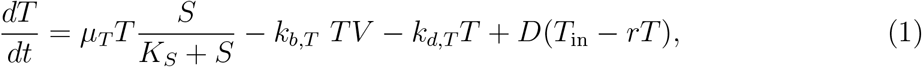

where *µ*_*T*_ is the growth kinetic constant for uninfected cells, *S* [nmol/mL] is the substrate concentration in the system, *K*_*S*_ is the Michaelis-Menten constant for substrate limitation to cell growth, *k*_*b,T*_ is the kinetic constant for viral binding to uninfected cells, *k*_*d,T*_ is the death kinetic constant for uninfected cells, *D* is the dilution rate of the system, *T*_in_ is the uninfected cells concentration in the feed, and *r* is the bleeding ratio. Table 1 clarifies the physical meaning of *D* and *r*. The terms on the right-hand side of Eq. 1 account for, from the left to the right: cell growth, viral infection, cell death, and inlet and outlet from the system. The balance of *i*(*t, τ*) [cell mL^−1^ hpi^−1^], the continuous distribution of the concentration of infected cells with respect to the infection age *τ*, is

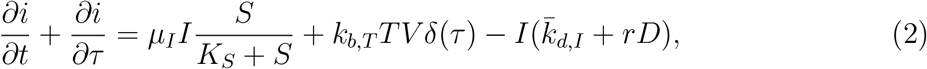

where *µ*_*I*_ is the growth kinetic constant of *i*,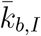 is the binding equivalent kinetic constant for infected cells,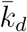 is the death equivalent kinetic constant for infected cells, and *δ*(·) is the Dirac delta function, which ensures that the contribution from newly infected cells is considered at the boundary (*τ* = 0). The terms in the right-hand side of Eq. 2 account for, from the left to the right: cell growth, infection of uninfected cells, cell death, and outlet from the system. For most viral species, infected cells undergo cell cycle arrest (*µ*_*I*_ = 0). The binding equivalent kinetic constant for infected cells depends on *τ*,

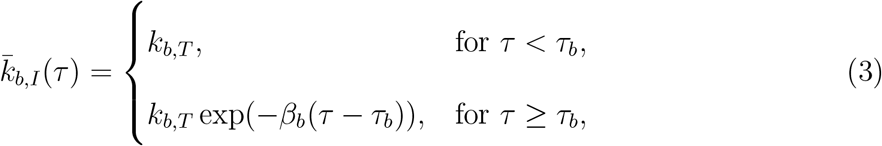

where *β*_*b*_ and *τ*_*b*_ are suitable coefficients that describe the viral binding downregulation with the progress of the infection age, which is experienced for most viral infections. ^47,48^ The death equivalent kinetic parameter depends on both the infection age and the viral genome copy number:^22^

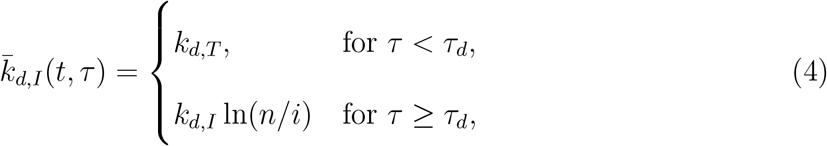

where *k*_*d,I*_ is the death kinetic constant for infected cells and *τ*_*d*_ is the infection age at which an increase of death rate is registered for infected cells, and *n/i* is the viral genome copy number in the nucleus of cells, as further discussed. Equation 4 establishes a dependence of the death rate of infected cells *i*(*t, τ*) from the intracellular level of viral genome. The balance of extracellular virions *V* (*t*) is

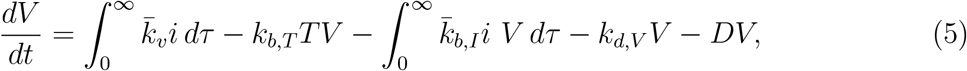

where *k*_*d,V*_ is the degradation kinetic constant for extracellular virions, and 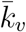 is the progeny release equivalent parameter. The terms on the right-hand side of Eq. 5 represent, from the left to the right: virion production from infected cells, viral infection of uninfected ells, viral infection of already-infected cells, degradation of extracellular virions, and outlet from the system. The progeny release per cell depends on the infection age:

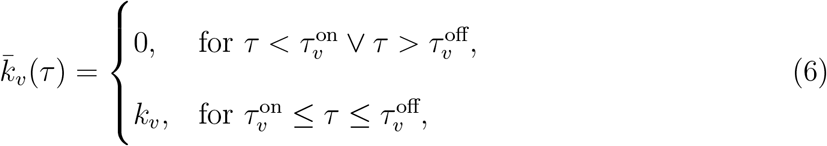

where *k*_*v*_ is the progeny release rate, and 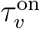 and 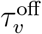define, respectively, the upper and lower bound of the infection age interval during which viral progeny is released. Equation 6 represents an effective way of lumping together several steps of the intracellular pathway that occur in between the infection of a cell and the onset of progeny release. For simplicity, the presence of only one substrate *S* is considered in the default implementation of the model.

The mass balance for *S*(*t*) is

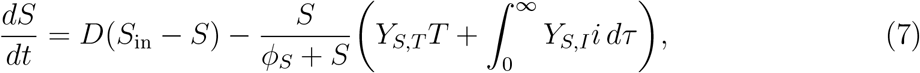

where *S*_in_ is the substrate concentration in the feed, *Y*_*S,T*_ is the specific substrate consumption for 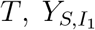 is the specific substrate consumption for 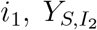 is the specific substrate consumption for *i*_2_, *Y*_*S,C*_ is the specific substrate consumption for *c*, and *ϕ*_*S*_ is a parameter describing the kinetics of substrate consumption at low substrate levels, which is fixed to *ϕ*_*S*_ = 0.01 nmol mL^−1^ in this work. A balance is also developed for tracking the concentration of nonviable cells *W* (*t*) in the system:

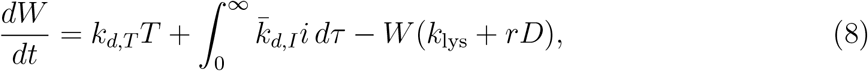

where *k*_lys_ is the lysis kinetic constant for nonviable cells. Species *W* considers all nonviable cells lumped together, independently if they were infected or not.

The model calculates the concentration of two intracellular species: *b*(*t, τ*), the virus bound to the surface of infected cells, and *n*(*t, τ*), the viral genome in the nucleus of a cell. Both *b*(*t, τ*) and *n*(*t, τ*) are computed with respect to the whole volume of the system, namely with unit of measurement [# mL^−1^ hpi^−1^]. This choice allows tracking the number of viruses uptaken by every cell more accurately, and also leads to greater numeric robustness. Notably, under the deterministic modeling assumptions followed here, cells infected at the same time instant will inherently present the same *b*(*t, τ*) and *n*(*t, τ*) profiles with the progress of time (until cell death). Hence, distributions *b*(*t, τ*) and *n*(*t, τ*) directly map to the distribution of cells with respect to the infection age *i*(*t, τ*). Normalization of *b*(*t, τ*) and *n*(*t, τ*) by *i*(*t, τ*) allows to convert the concentration of the intracellular species into distributions expressed in a per cell basis [# */*cell]. The model assumes irreversible viral attachment, hence a cell is considered infected as soon as a virus bounds to its surface. Species *b* can represent viruses either bound to receptors or directly attached to the cell membrane. The model parameters that govern viral binding inherently account for the trade-off between binding and unbinding rates. The balance for *b*(*t, τ*) is

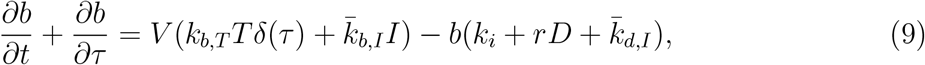

where the terms on the right-hand side of the equation represent, from the left to the right: viral binding to uninfected cells, viral binding to infected cells, internalization of surface-attached viruses through kinetic constant *k*_*i*_, outlet from the system, and death of infected cells. The last term is included to remove from the mass balance the amount of virus bound to the surface of infected cells that die, since *b* represents the cumulative amount in the system of virus bound to (viable) infected cells. The model considers trafficking to nucleus as a single lumped step. This simplification is introduced to reduce the computational burden of the model, especially for its two-dimensional implementation. This modeling choice does not significantly degrade the model predictive performance, as long as *k*_*i*_ is taken as the kinetic parameter for the rate determining step within the pathway of trafficking to nucleus.

The balance for *n*(*t, τ*) is

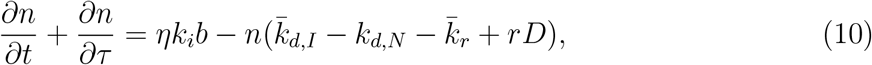

where the terms on the right-hand side represent, from left to right: trafficking to nucleus, contribution due to cell death (as for *b*), degradation of viral genome in the nucleus (with kinetic parameter *k*_*d,N*_), viral replication, and outlet of infected cells from the system. Parameter *η* represents the fraction of internalized virus that reaches the nucleus, and also accounts for rerouting to lysosomes. The effective replication kinetic parameter 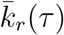 is calculated as

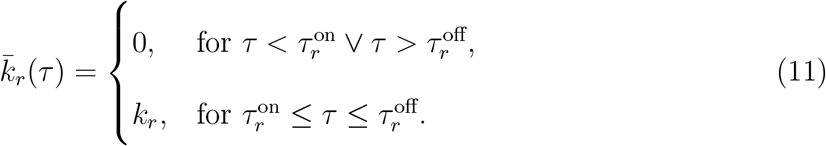

where *k*_*r*_ is the viral genome replication kinetic constant, and 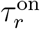 and 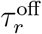 define, respectively, the upper and lower bound of the infection age interval during which viral amplification occurs. Equation 11 represents an efficient way of lumping together several steps of the intracellular pathway for viral amplification. Notably, 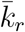 should be set equal to the cell growth rate for viruses that integrate into the host genome. The set of Eqs. 1, 2, 5, 7–10 represents the model for systems with one viral species. The list of model parameters is summarized in Table S3.

### Numerics

A decoupled integration-reallocation numerical scheme is introduced in this work. The model equations are integrated with the method of lines, by discretizing the PDEs into ODEs along the infection age coordinate(s) (Fig. 2). During integration, infection ages are assumed not to vary. Separately, a reallocation routine accounts for the increase of infection age, as further discussed. The numerical scheme is first described in detail for the simplified model for systems with one viral species, outlined in the previous section. The infection age *τ* is discretized into *L* nodes. All nodes have size Δ*τ*, except for node *L*, which corresponds to the interval [*τ*_max_, ∞) in the original space of the continuous variable *τ*. The maximum infection age *τ*_max_, arbitrarily selected, is an infection age associated to very low cell viability as well as negligible recombinant product expression and progeny release. Mesh nodes *l* = 1, 2, 3, …, *L* have infection age *τ* ^*l*^ = 0, Δ*τ*, 2Δ*τ*, …, *τ*_max_. The distributed states *b*(*t, τ*), *i*(*t, τ*), and *n*(*t, τ*) are discretized with respect to *τ* into the sets of *L* states denoted as, respectively, *B*(*t, τ* ^*l*^) = *B*^*l*^, *I*(*t, τ* ^*l*^) = *I*^*l*^, and *N* (*t, τ* ^*l*^) = *N* ^*l*^, for *l* = 1, 2, …, *L*. The model PDEs (Eqs. 2, 9–10) are converted into the sets of ODEs:

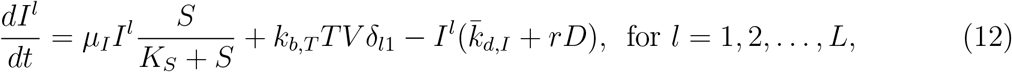

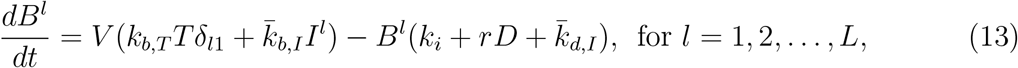

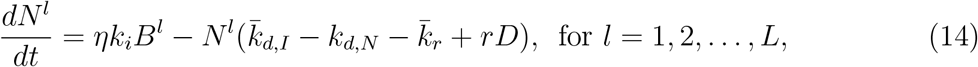

where *δ*_*l*1_ is the Kronecker delta function. Equations 12–14 do not account for the 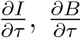 and 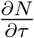 contributions in Eqs. 2, 9–10, but present otherwise the same contributions of the original PDEs. The integrals in Eqs. 5, 7, 8 are converted into sums across all mesh nodes. The ODE/PDE system of Eqs. 1, 2, 5, 7–10 becomes the system of ODEs of Eqs. 1, 5, 7, 8, 12–14, which are integrated with a third-order Runge-Kutta scheme with adaptive time stepping. At every integration step *i*, the cumulative sum of the integration steps (*s*_*i*_) is updated with the *i*th integration step *h*_*i*_ as

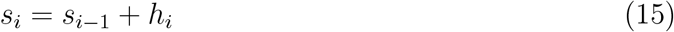

where *s*_0_ is initialized to 0. At the end of each integration step *i*, if the condition

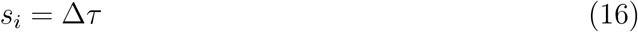

is met, then *I*^*l*^, *B*^*l*^, and *N* ^*l*^ (for *l* = 1, 2, …, *L*) are reallocated along the mesh to account for the increase of infection age, with the following procedure. Defining the value of the states *I*^*l*^, *B*^*l*^, and *N* ^*l*^ in every mesh node at the end of integration step *i* as, respectively, *I*^*l,i*,fin^, *B*^*l,i*,fin^, and *N* ^*l,i*,fin^, the corresponding initial value of the states for integration step *i* + 1 (respectively, *I*^*l,i*+1,in^, *B*^*l,i*+1,in^, and *N* ^*l,i*+1,in^) is calculated as

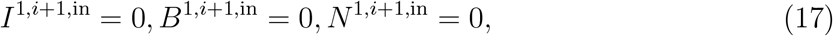

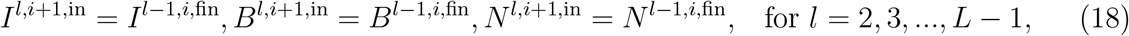

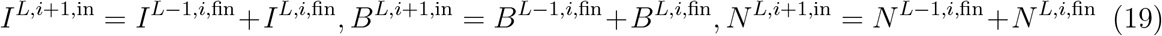

The integration step counter *i* is then re-initialized to 0. At the end of every integration step *i* in which Condition 16 is not met, *I*^*l,i*+1,in^, *B*^*l,i*+1,in^, and *N* ^*l,i*+1,in^ correspond, for every mesh node *l*, to, respectively, *I*^*l,i*,fin^, *B*^*l,i*,fin^, and *N* ^*l,i*,fin^. The described integration-reallocation steps are iteratively repeated during simulation.

The model for systems with two viruses is solved with an analogous approach. The continuous variables *τ*_1_ and *τ*_2_ are both discretized into *L* mesh nodes (respectively, 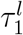 and 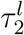 for *l* = 1, 2, …, *L*). All nodes have size Δ*τ*, except for 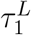 and 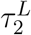, which correspond to the interval [*τ*_max_, ∞) in the original space of *τ*_1_ and *τ*_2_. The distributed states are discretized with respect to *τ*_1_ and *τ*_2_, as shown in Fig. 2. To simplify the notation, the 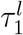 and 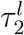 notation is dropped and, instead, it is postulated that *τ*_1_ and *τ*_2_ can only assume discrete values when they refer to discrete distributions (denoted by uppercase letters). Accordingly, the integrals over the infection age(s) in the model equations (Eqs. S9–S13) are converted into sums across all mesh nodes. The PDEs corresponding to distributed states (Eqs. S2–S3, S6, S14–S21) are converted into sets of ODEs, as outlined for the model for systems with one virus. The ODEs resulting from the discretization are solved, together with the other ODEs of the model (Eqs. S1, S9–S13), with an integration-reallocation approach. The reallocation for states distributed with respect to only *τ*_1_ or *τ*_2_ (i.e., *I*_1_, *I*_2_, 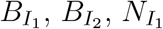, and 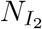) is carried out with a procedure analogous to Eqs. 17–19. States distributed with respect to both *τ*_1_ and *τ*_2_ (i.e., 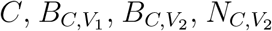 and 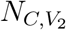) are reallocated along both *τ*_1_ and *τ*_2_ simultaneously, resulting in diagonal movements along the mesh (Fig. 2). Boundary conditions analogous to Eqs. 17 and 19 are imposed also for the states distributed to both infection ages. Finally, the viral binding downregulation experienced for most viral infections (Eq. 3) is exploited to reduce the computational burden. A parameter *τ*_*b*,max_ ≤ *τ*_max_ is defined, based on the viral binding decay profile with respect to the infection age for the considered viral system. In all equations distributed with respect to one or more infection ages, the contributions from viral binding are not computed in mesh nodes where the infection age is equal or greater than *τ*_*b*,max_. Further, the nodes of the two-dimensional infection age mesh in which |*τ*_1_ − *τ*_2_| ≥ *τ*_*b*,max_ cannot be accessed by any species. The corresponding states are set to 0 for every value of *t*, and the associated ODEs are not computed. This approach for reducing the number of equations of the model is not used if DIPs are present in the system, since, typically, DIP-infected cells can always be re-infected, independently from the infection age with respect to DIPs.

### Simulation

The results presented here were generated with a MATLAB implementation of the model and of the numerics introduced in this work. The complete simulation settings are summarized in Table S6 for all case studies. The model for systems with one virus is used in Case studies 1–2, while the model for systems with two viruses is used in Case studies 3–4. In Case study 4, two additional ODEs are solved to compute the mass balance of viral genome 1 and 2 in nonviable cells, to generate the results reported in Fig. S5. A third additional ODE is solved in Case study 4 for tracking the total concentration of extracellular DIPs (Fig. 9), namely the sum of infectious PFUs, considered by the default implementation of the model) and non-infectious DIP virions. For this equation, following literature findings,^44^the ratio between total released DIPs and infectious released DIPs is approximated to the value of 300. Regarding the model parameters, Case studies 1–3 are directly carried out using validated parameters from the literature.^22^In Case study 4, several parameters are fixed based on the intrinsic characteristics of STV/DIP systems or directly obtained from the literature (Table S5). The remaining 11 model parameters are estimated through experimental data reported by Rudiger et al. ^21^Parameter estimation is carried out with maximum likelihood estimation^49^on measurements of STV and DIP titer (Fig. 9), intracellular STV and DIP viral genome copy number (Fig. S5), and VCD (Fig. S6) for 8 out of the 12 experiments reported by Rudiger et al. ^21^Figure 9 clarifies the division of the experiments into training (used for parameter estimation) and validation datasets. Further, the extracellular and intracellular STV concentration measurements discussed here refer to the concentrations of full-length influenza A segment 1 RNA reported by Rudiger et al. Although Rudiger et al. also provide measurements for the concentration of segment 5, full-length segment 1 is selected as a proxy for the STV concentration, since DI244 present its genomic deletion in segment 1.

## Abbreviations

AIC: Akaike information criterion
BEVS: baculovirus expression vector system
DIP: defective interfering particle
hpi: hours post infection
MDCK: Madin-Darby canine kidney
MOI: multiplicity of infection
ODE: ordinary differential equation
PB2: polymerase basic protein 2
PFU: plaque-forming unit
PDE: partial differential equation
RSS: residual sum of squares
rAAV: recombinant adeno-associated virus
STV: standard virus
TOI: time of infection
VCD: viable cell density
VLP: virus-like particle
vg: viral genome

## Acknowledgement

This work was done in Cambridge, MA, USA. This work was supported by the U.S. Food and Drug Administration through Contract no. 75F40121C00131.

## Code Availability

Once the manuscript is accepted, an implementation of the simulator presented in this work will be downloadable at https://github.com/francescodestro/vitraPro.

## Supporting Information Available

The following files are available free of charge.

- Supporting Information: Additional model benchmarking, list of model parameters and simulation inputs, discussion of model limitations, additional information on Case Study 4, and supplementary methods

