## Supporting Information for "Efficient Simulation of Viral Transduction and Propagation for Biomanufacturing"

Table S1: Computational time: average among 10 runs of the same simulation. Simulations are performed in MATLAB on an Intel Core i9-13950HX CPU@2.20GHz processor.

| Case study | Exp. | MOI | Process duration | Clock time |
|---|---|---|---|---|
| 1 | 1 | $10^{-3}$ | 10 d | 0.72 s |
| | 2 | $10^{-1}$ | 10 d | 0.60 s |
| | 3 | 1 | 10 d | 0.74 s |
| 2 | 1 | $10^{-3}$ | 10 d | 1.62 s |
| | 2 | $10^{-1}$ | 10 d | 1.61 s |
| | 3 | 1 | 10 d | 1.53 s |
| 3 | 1 | – | 10 d | 166.86 s |
| 4 | 1 | STV: $10^{-3}$, DIP: 0 | 35.75 h | 4.84 s |
| | 2 | STV: $10^{-3}$, DIP: $10^{-3}$ | 35.75 h | 37.03 s |
| | 3 | STV: $10^{-3}$, DIP: 3 | 64.75 h | 30.35 s |
| | 4 | STV: $10^{-3}$, DIP: 30 | 40.75 h | 5.39 s |
| | 5 | STV: 3, DIP: 0 | 35.75 h | 8.22 s |
| | 6 | STV: 3, DIP: $10^{-3}$ | 35.75 h | 53.23 s |
| | 7 | STV: 3, DIP: 3 | 35.75 h | 14.70 s |
| | 8 | STV: 3, DIP: 30 | 35.75 h | 23.57 s |
| | 9 | STV: 30, DIP: 0 | 35.75 h | 10.79 s |
| | 10 | STV: 30, DIP: $10^{-3}$ | 35.75 h | 15.29 s |
| | 11 | STV: 30, DIP: 3 | 35.75 h | 12.90 s |
| | 12 | STV: 30, DIP: 30 | 35.75 h | 21.58 s |

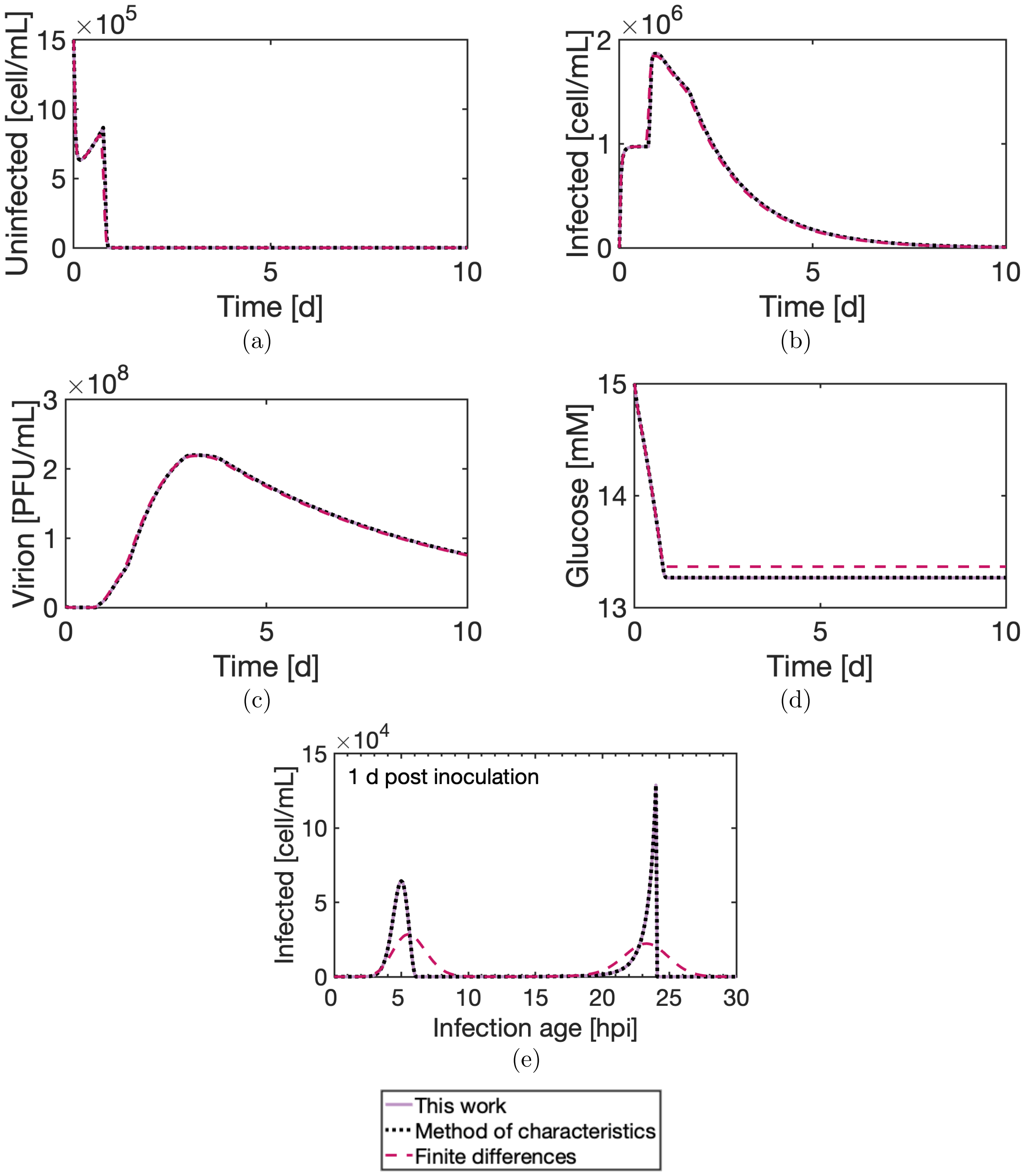


Figure S1: Comparison of numerical methods for Case study 1 (scenario with inoculation at MOI = 1 PFU per cell). The solution derived with the numerical method introduced in this work is benchmarked against those derived from the method of characteristics and finite differences: (a) uninfected cells concentration, (b) infected cells concentration, (c) virion concentration, (d) glucose concentration, (e) infection age distribution of the infected cells 1 day after inoculation. The distributions in (e) appear continuous, but are discrete frequency distribution with bin size $\Delta\tau = 0.1$ hpi.

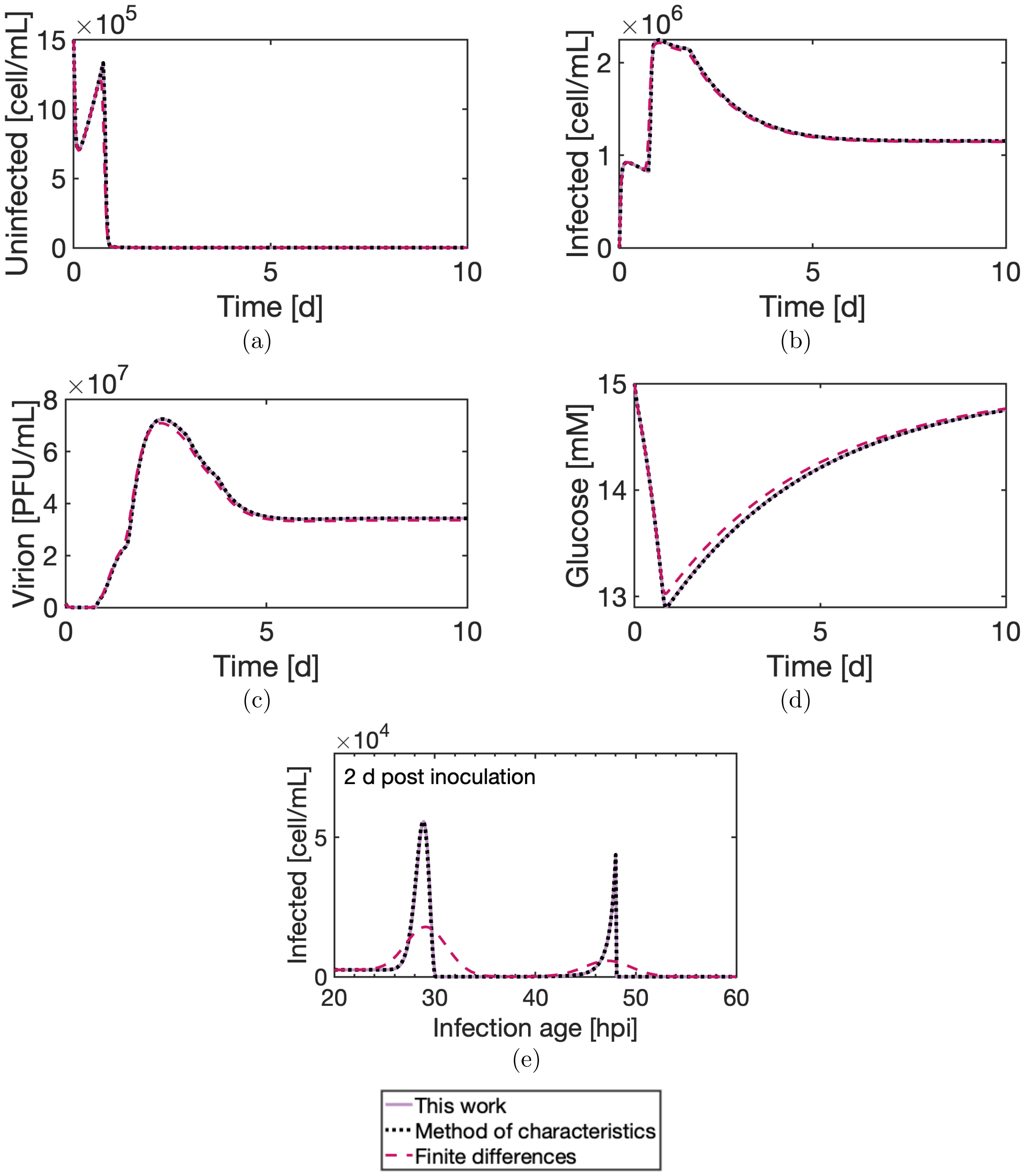


Figure S2: Comparison of numerical methods for Case study 2 (scenario with inoculation at MOI = 1 PFU per cell). The solution derived with the numerical method introduced in this work is benchmarked against those derived from the method of characteristics and finite differences: (a) uninfected cells concentration, (b) infected cells concentration, (c) virion concentration, (d) glucose concentration, (e) infection age distribution of the infected cells 2 day after inoculation. The distributions in (e) appear continuous, but are discrete frequency distribution with bin size $\Delta\tau = 0.1$ hpi.

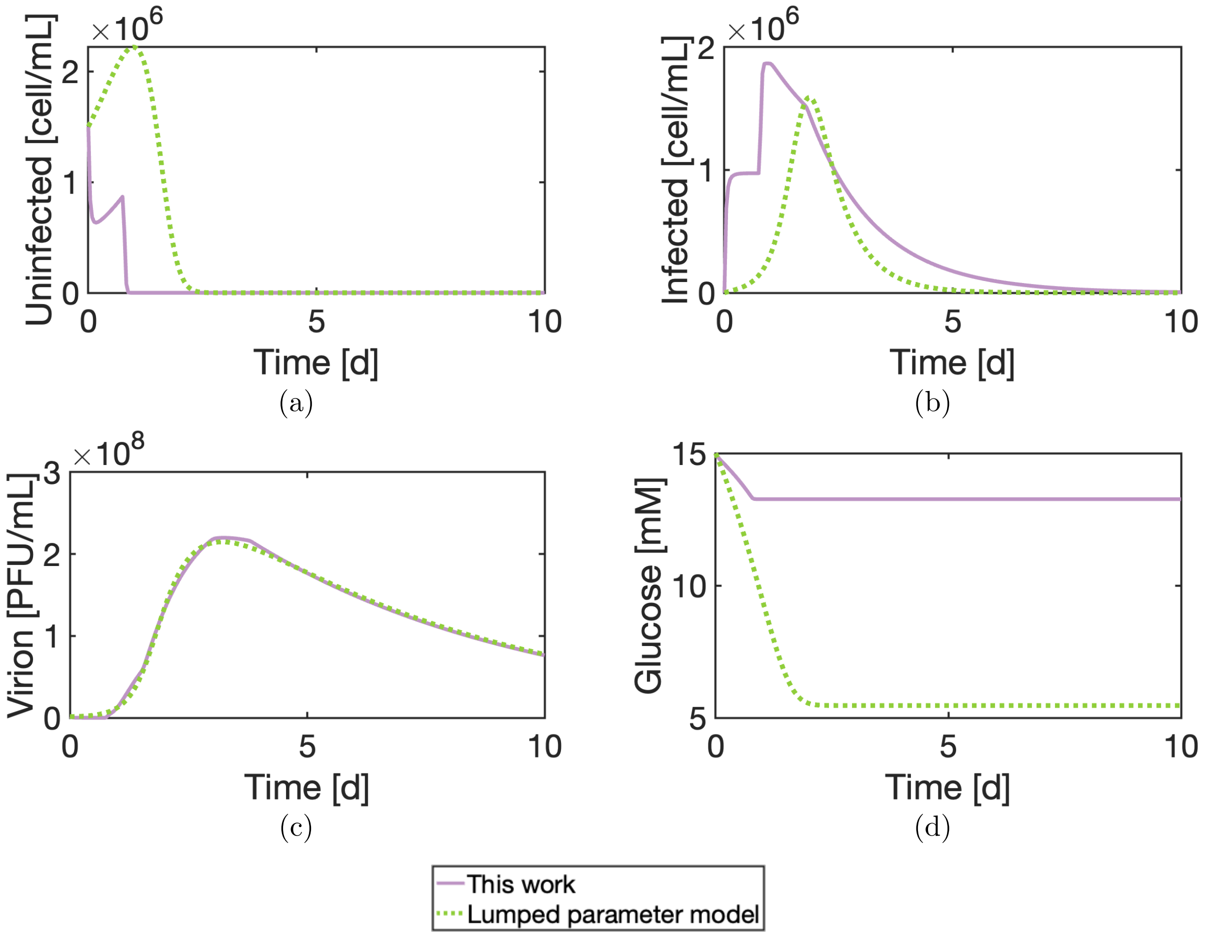


Figure S3: Comparison with the lumped parameter model: Case study 1 (scenario with inoculation at MOI = 1 PFU per cell). (a) uninfected cells concentration, (b) infected cells concentration, (c) virion concentration, and (d) glucose concentration. Additional information on this comparison is provided in the Supplementary Methods.

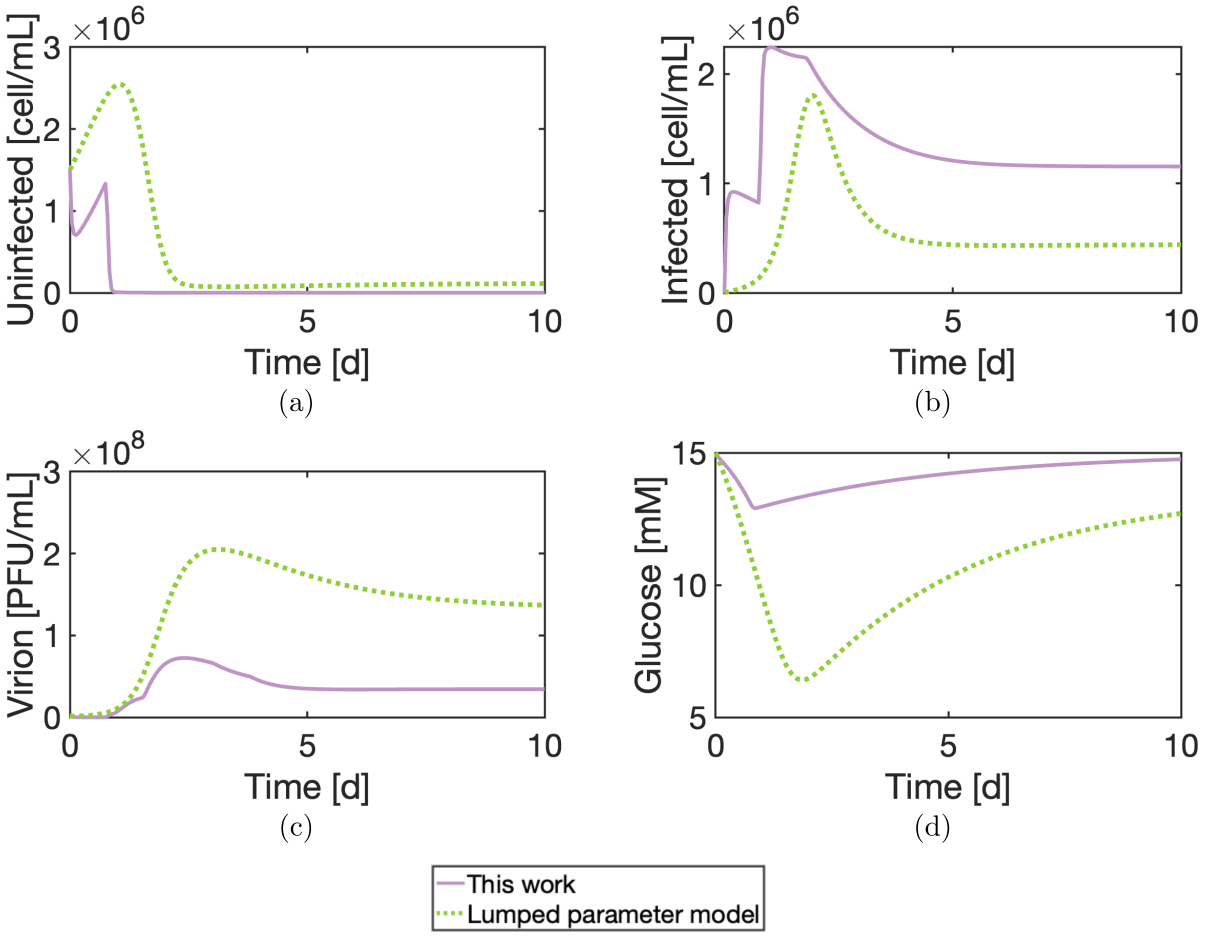


Figure S4: Comparison with the lumped parameter model: Case study 2 (scenario with inoculation at MOI = 1 PFU per cell). (a) uninfected cells concentration, (b) infected cells concentration, (c) virion concentration, and (d) glucose concentration. Additional information on this comparison is provided in the Supplementary Methods.

Table S2: Model parameters for systems with two viruses.

| Symbol | UOM | Description |
|---|---|---|
| Cell growth and substrate consumption | | |
| $\mu_k$ | $h^{-1}$ | Growth kinetic constant for cells $k = \{T, I_1, I_2, C\}$ |
| $K_S$ | nmol $mL^{-1}$ | Michaelis-Menten constant for substrate limitation to cell growth |
| $Y_{S,k}$ | nmol $cell^{-1}$ | Substrate consumption for cells $k = \{T, I_1, I_2, C\}$ |
| Binding | | |
| $k_{b,T,V_j}$ | mL $cell^{-1}$ $h^{-1}$ | Kinetic constant for binding of virus $j = \{1, 2\}$ to $T$ |
| $\tau_{b,k}$ | hpi | Onset of decay of viral binding rate for cells $k = \{I_1, I_2\}$ |
| $\beta_{b,k}$ | [–] | Coefficient for decay of viral binding rate for cells $k = \{I_1, I_2\}$ |
| $\tau_{b,C,V_j}$ | hpi | Onset of decay of binding rate of virus $j = \{1, 2\}$ to coinfected cells |
| $\beta_{b,C,V_j}$ | [–] | Coefficient for decay of binding rate of virus $j = \{1, 2\}$ to coinfected cells |
| Cell death | | |
| $f_{d,C,V_j}$ | [–] | Contribution of virus $j = \{1, 2\}$ to coinfected cell death rate |
| $k_{d,k}$ | $h^{-1}$ | Death kinetic constant for cells $k = \{T, I_1, I_2, C\}$ |
| $k_{lys}$ | $h^{-1}$ | Lysis rate of nonviable cells |
| $\tau_{d,k}$ | hpi | Onset of death rate increase for cells $k = \{I_1, I_2, C\}$ |
| Viral degradation | | |
| $k_{d,V_j}$ | $h^{-1}$ | Degradation kinetic constant for extracellular virus $j = \{1, 2\}$ |
| $k_{d,N_j}$ | $h^{-1}$ | Degradation kinetic constant for nuclear viral genome of virus $j = \{1, 2\}$ |
| Virus internalization | | |
| $k_{i,k}$ | $h^{-1}$ | Trafficking to nucleus kinetic constant for $k = \{I_1, I_2\}$ |
| $k_{i,C,V_j}$ | $h^{-1}$ | Trafficking to nucleus kinetic constant for virus $j = \{1, 2\}$ in coinfected cells |
| $\eta_j$ | [–] | Fraction of virus $j = \{1, 2\}$ internalized in $I_i$ that reaches the nucleus |
| $\eta_{i,C,V_j}$ | [–] | Fraction of virus $j = \{1, 2\}$ internalized in $C$ that reaches the nucleus |
| Viral replication | | |
| $k_{r,k}$ | $h^{-1}$ | Replication kinetic constant of viral genome in $k = \{I_1, I_2\}$ |
| $k_{r,C,V_j}$ | $h^{-1}$ | Replication kinetic constant of viral genome $j = \{1, 2\}$ in $C$ |
| $\tau_{r,k}^{on}$ | hpi | Onset of viral genome replication replication in $k = \{I_1, I_2\}$ |
| $\tau_{r,k}^{off}$ | hpi | End of viral genome replication in $k = \{I_1, I_2\}$ |
| $\tau_{r,C,V_j}^{on}$ | hpi | Onset of viral genome $j = \{1, 2\}$ replication in $C$ |
| $\tau_{r,C,V_j}^{off}$ | hpi | End of viral genome $j = \{1, 2\}$ replication in $C$ |

| Symbol | UOM | Description |
|---|---|---|
| Progeny release | | |
| $k_{v,k}$ | PFU cell$^{-1}$ h$^{-1}$ | Progeny release rate for $k = \{I_1, I_2, C\}$ |
| $\tau_{v,k}^{\text{on}}$ | hpi | Onset of progeny release for $k = \{I_1, I_2, C\}$ |
| $\tau_{v,k}^{\text{off}}$ | hpi | End of progeny release for $k = \{I_1, I_2, C\}$ |
| $\phi_{I_2,V_1}$ | [–] | Coefficient for random generation of $V_1$ from $I_2$ |
| $\phi_{I_1,V_2}$ | [–] | Coefficient for random generation of $V_2$ from $I_1$ |

Table S3: Model parameters for systems with only one virus. The parameters used for simulating the BEVS in Case studies 1 and 2 are reported.

| Symbol | UOM | Description | Value[S1–S3] |
|---|---|---|---|
| Cell growth and substrate consumption | | | |
| $\mu_T$ | h$^{-1}$ | Growth kinetic constant for uninfected cells | 0.028 |
| $\mu_I$ | h$^{-1}$ | Growth kinetic constant for infected cells | 0 |
| $K_S$ | nmol mL$^{-1}$ | Michaelis-Menten constant for substrate limitation to cell growth | $1.3\times10^{3}$ |
| $Y_{S,T}$ | nmol cell$^{-1}$ h$^{-1}$ | Substrate consumption for infected cells | $1.2\times10^{-4}$ |
| $Y_{S,I}$ | nmol cell$^{-1}$ h$^{-1}$ | Substrate consumption for infected cell | $0^{*}$ |
| Binding | | | |
| $k_{b,T}$ | mL cell$^{-1}$ h$^{-1}$ | Binding kinetic constant | $6.3\times10^{-7}$ |
| $\tau_b$ | hpi | Onset of decay of viral binding rate for infected cells | 1.80 |
| $\beta_b$ | [–] | Coefficient for decay of viral binding rate for infected cells | 0.5 |
| Cell death | | | |
| $k_{d,T}$ | h$^{-1}$ | Death kinetic constant for uninfected cells | $8\times10^{-5}$ |
| $k_{d,I}$ | h$^{-1}$ | Death kinetic constant for infected cells | $2.9\times10^{-3}$ |
| $k_{lys}$ | h$^{-1}$ | Lysis rate of nonviable cells | $0^{\dagger}$ |
| $\tau_d$ | hpi | Onset of death rate increase for infected cells | 24 |
| Viral degradation | | | |
| $k_{d,V}$ | h$^{-1}$ | Degradation kinetic constant for extracellular virus | $7\times10^{-3}$ |
| $k_{d,N}$ | h$^{-1}$ | Degradation kinetic constant for nuclear viral genome | 0 |
| Virus internalization | | | |

*Substrate consumption for infected cells was neglected, although it can be non-null in early infection stages.[S3]

†Cell lysis not considered in simulations for Case studies 1–2.

| Symbol | UOM | Description | Value[S1–S3] |
|---|---|---|---|
| $k_i$ | $h^{-1}$ | Virus internalization kinetic constant | 0.6 |
| $\eta$ | [–] | Fraction of internalized virus that reaches the nucleus | 0.50 |
| Viral replication | | | |
| $k_r$ | $h^{-1}$ | Viral genome replication kinetic constant | 0.732 |
| $\tau_r^{\mathrm{on}}$ | hpi | Onset of viral genome replication replication in infected cells | 6 |
| $\tau_r^{\mathrm{off}}$ | hpi | End of viral genome replication in infected cells | 18 |
| Progeny release | | | |
| $k_v$ | PFU cell$^{-1}$ h$^{-1}$ | Progeny release rate for infected cells | 5 |
| $\tau_{v,k}^{\mathrm{on}}$ | hpi | Onset of progeny release for infected cells | 18 |
| $\tau_{v,k}^{\mathrm{off}}$ | hpi | End of progeny release for infected cells | 72 |

Table S4: Parameters used for model simulation in Case study 3 (co-transduction from two recombinant baculoviruses). The description of the parameters is reported in Table S2. The reported equalities among parameters are typical of systems characterized by the presence of two recombinant viruses of the same type, differing only in the recombinant cassette.

| Symbol | UOM | Value[S1–S3] |
|---|---|---|
| Cell growth and substrate consumption | | |
| $\mu_T$ | $h^{-1}$ | 0.028 |
| $\mu_{I_1} = \mu_{I_2} = \mu_C$ | $h^{-1}$ | 0 |
| $K_S$ | nmol $mL^{-1}$ | $1.3\times10^{3}$ |
| $Y_{S,T}$ | nmol $cell^{-1}$ $h^{-1}$ | $1.2\times10^{-4}$ |
| $Y_{S,I_1} = Y_{S,I_2} = Y_{S,C}$ | nmol $cell^{-1}$ $h^{-1}$ | $0^{\dagger}$ |
| Binding | | |
| $k_{b,T,V_1} = k_{b,T,V_2}$ | mL $cell^{-1}$ $h^{-1}$ | $6.3\times10^{-7}$ |
| $\tau_{b,I_1} = \tau_{b,I_2} = \tau_{b,C,V_1} = \tau_{b,C,V_2}$ | hpi | 1.80 |
| $\beta_{b,I_1} = \beta_{b,I_2} = \beta_{b,C,V_1} = \beta_{b,C,V_2}$ | [–] | 0.5 |
| Cell death | | |
| $k_{d,T}$ | $h^{-1}$ | $8\times10^{-5}$ |
| $k_{d,I_1} = k_{d,I_2} = k_{d,C}$ | $h^{-1}$ | $2.9\times10^{-3}$ |
| $\tau_{d,I_1} = \tau_{d,I_2} = \tau_{d,C}$ | hpi | 24 |
| $f_{d,C,V_1} = f_{d,C,V_2}$ | [–] | 1 |
| $k_{lys}$ | $h^{-1}$ | $0^{*}$ |
| Viral degradation | | |
| $k_{d,V_1} = k_{d,V_2}$ | $h^{-1}$ | $7\times10^{-3}$ |
| $k_{d,N_1} = k_{d,N_2}$ | $h^{-1}$ | 0 |
| Virus internalization | | |
| $k_{i,I_1} = k_{i,I_2} = k_{i,C,V_1} = k_{i,C,V_2}$ | $h^{-1}$ | 0.6 |
| $\eta_{i,I_1} = \eta_{i,I_2} = \eta_{i,C,V_1} = \eta_{i,C,V_2}$ | [–] | 0.50 |
| Viral replication | | |
| $k_{r,I_1} = k_{r,I_2} = k_{r,C,V_1} = k_{r,C,V_2}$ | $h^{-1}$ | 0.732 |
| $\tau^{on}_{r,I_1} = \tau^{on}_{r,I_2} = \tau^{on}_{r,C,V_1} = \tau^{on}_{r,C,V_2}$ | hpi | 6 |
| $\tau^{off}_{r,I_1} = \tau^{off}_{r,I_2} = \tau^{off}_{r,C,V_1} = \tau^{off}_{r,C,V_2}$ | hpi | 18 |
| Progeny release | | |
| $k_{v,I_1} = k_{v,I_2} = k_{v,C}$ | PFU $cell^{-1}$ $h^{-1}$ | 5 |
| $\tau^{on}_{v,I_1} = \tau^{on}_{v,I_2} = \tau^{on}_{v,C}$ | hpi | 18 |
| $\tau^{off}_{v,I_1} = \tau^{off}_{v,I_2} = \tau^{off}_{v,C}$ | hpi | 72 |
| $\phi_{I_2,V_1} = \phi_{I_1,V_2}$ | [–] | $0^{\ddagger}$ |

†Substrate consumption for infected cells was neglected, although it can be non-null in early infection stages.[S3]

†Generally valid for systems with two STVs

Table S5: Parameters used for model simulation in Case study 4 (influenza propagation in presence of DIPs). The reported equalities among parameters are typical of systems characterized by the presence of one STV and one DIP of the same type of virus. The full description of the parameters is reported in Table S2. STV: virus 1; DIP: virus 2.

| Symbol | UOM | Value | Details |
|---|---|---|---|
| Cell growth and substrate consumption | | | |
| $\mu_T = \mu_{I_2}$ | $h^{-1}$ | 0.02 | Fixed based on doubling time of cell/media system[S4] |
| $\mu_{I_1} = \mu_C$ | $h^{-1}$ | 0 | Fixed† |
| $K_S$ | nmol $mL^{-1}$ | – | Substrate not limiting in considered cell/media system[S4] |
| $Y_{S,T} = Y_{S,I_2}$ | nmol $cell^{-1}h^{-1}$ | – | Substrate not limiting in considered cell/media system[S4] |
| $Y_{S,I_1} = Y_{S,C}$ | nmol $cell^{-1}h^{-1}$ | – | Substrate not limiting in considered cell/media system[S4] |
| Binding | | | |
| $k_{b,T,V_1} = k_{b,T,V_2}$ | mL $cell^{-1}$ $h^{-1}$ | $1.5\times10^{-6}$ | Estimated from Figs. 9, S5, S6 |
| $\tau_{b,I_1} = \tau_{b,C,V_1} = \tau_{b,C,V_2}$ | hpi | 1.5 | Fixed considering superinfection limitation dynamics reported in literature[S5] |
| $\beta_{b,I_1} = \beta_{b,C,V_1} = \beta_{b,C,V_2}$ | [–] | 10 | Fixed to high value, compatible with fast decay of binding rate |
| $\tau_{b,I_2}$ | [–] | 15 | Fixed to high value† |
| $\beta_{b,I_2}$ | [–] | 100 | Fixed to high value,† compatible with fast decay of binding rate |
| Cell death | | | |
| $f_{d,C,V_1}$ | [–] | 1 | Fixed† |
| $f_{d,C,V_2}$ | [–] | 0 | Fixed† |
| $k_{d,T} = k_{d,I_2}$ | $h^{-1}$ | $8.6\times10^{-3}$ | Estimated from Figs. 9, S5, S6 |
| $k_{d,I_1} = k_{d,C}$ | $h^{-1}$ | $3\times10^{-2}$ | Estimated from Figs. 9, S5, S6 |
| $k_{lys}$ | $h^{-1}$ | 0.15 | Estimated from Figs. 9, S5, S6 |
| $\tau_{d,I_1} = \tau_{d,C}$ | hpi | 6.5 | Estimated from Figs. 9, S5, S6 |
| Viral degradation | | | |
| $k_{d,V_1} = k_{d,V_2}$ | $h^{-1}$ | 0.17 | Estimated from Figs. 9, S5, S6 |
| $k_{d,N_1} = k_{d,N_2}$ | $h^{-1}$ | 0 | Fixed |
| Virus internalization | | | |
| $k_{i,I_1} = k_{i,I_2} = k_{i,C,V_1} = k_{i,C,V_2}$ | $h^{-1}$ | 3 | Kinetic rate of limiting step for trafficking to nucleus[S6] |
| $\eta_{i,I_1} = \eta_{i,I_2} = \eta_{i,C,V_1} = \eta_{i,C,V_2}$ | [–] | 0.5 | From literature[S6] |

†Generally valid for STV/DIP systems

| Symbol | UOM | Value | Details |
|---|---|---|---|
| Viral replication | | | |
| $k_{r,I_1} = k_{r,C,V_1}$ | $h^{-1}$ | 1.9228 | Estimated from Figs. 9, S5, S6 |
| $k_{r,I_2} = 0$ | $h^{-1}$ | 0 | Fixed† |
| $k_{r,C,V_2}$ | $h^{-1}$ | 2.3067 | Estimated from Figs. 9, S5, S6 |
| $\tau^{\mathrm{on}}_{r,I_1} = \tau^{\mathrm{on}}_{r,C,V_1} = \tau^{\mathrm{on}}_{r,C,V_2}$ | hpi | 2.55 | Estimated from Figs. 9, S5, S6 |
| $\tau^{\mathrm{off}}_{r,I_1} = \tau^{\mathrm{off}}_{r,C,V_1} = \tau^{\mathrm{off}}_{r,C,V_2}$ | hpi | 7.35 | Estimated from Figs. 9, S5, S6 |
| Progeny release | | | |
| $k_{v,I_1} = k_{v,C}$ | PFU $cell^{-1}$ $h^{-1}$ | 20 | Estimated from Figs. 9, S5, S6 |
| $k_{v,I_2} = 0$ | PFU $cell^{-1}$ $h^{-1}$ | | Fixed† |
| $\tau^{\mathrm{on}}_{v,I_1} = \tau^{\mathrm{on}}_{v,C}$ | hpi | 6.5 | From literature[S7] |
| $\tau^{\mathrm{off}}_{v,I_1} = \tau^{\mathrm{off}}_{v,C}$ | hpi | – | Cell death |
| $\phi_{I_2,V_1}$ | [–] | 0 | Fixed (no STV production from DIP-infected cells) |
| $\phi_{I_1,V_2}$ | [–] | 0 | Fixed (random DIP generation not important in considered dataset, since DIPs are inoculated) |

Table S6: Simulation inputs for Case studies 1–4. j={1,2}.

| | | Case study | | | |
|---|---|---|---|---|---|
| | UOM | 1 | 2 | 3 | 4 |
| $D$ | $h^{-1}$ | 0 | 0.01 | 0.01 | 0 |
| $r$ | [–] | – | 1 | 1 | – |
| $S_{\text{in}}$ | nmol $mL^{-1}$ | – | $15\times10^3$ | $15\times10^3$ | – |
| $T_{in}$ | cell $mL^{-1}$ | – | $3\times10^6$ | $3\times10^6$ | – |
| $B_{I_j}(0,\tau_j)$ | virus $mL^{-1}$ | $0,\ \forall j,\tau_j$ | $0,\ \forall j,\tau_j$ | $0,\ \forall j,\tau_j$ | $0,\ \forall j,\tau_j$ |
| $B_{0,V_j}(0,\tau_1,\tau_2)$ | virus $mL^{-1}$ | $0,\ \forall j,\tau_1,\tau_2$ | $0,\ \forall j,\tau_1,\tau_2$ | $0,\ \forall j,\tau_1,\tau_2$ | $0,\ \forall j,\tau_1,\tau_2$ |
| $C(t,\tau_1,\tau_2)$ | cell $mL^{-1}$ | $0,\ \forall\tau_1,\tau_2$ | $0,\ \forall\tau_1,\tau_2$ | $0,\ \forall\tau_1,\tau_2$ | $0,\ \forall\tau_1,\tau_2$ |
| $I_j(0,\tau_j)$ | cell $mL^{-1}$ | $0,\ \forall j,\tau_j$ | $0,\ \forall j,\tau_j$ | $3\times10^5$ for $j$=1,$\tau_1$=40 hpi;<br>$1.5\times10^5$ for $j$=2,$\tau_2$=40 hpi;<br>0 otherwise | $0,\ \forall j,\tau_j$; |
| $N_{I_j}(0,\tau_j)$ | vg $mL^{-1}$ | $0,\ \forall\tau_1,\tau_2$ | $0,\ \forall\tau_1,\tau_2$ | $1.5\times10^{10}$ for $j$=1,$\tau_1$=40 hpi;<br>$7.5\times10^9$ for $j$=2,$\tau_2$=40 hpi;<br>0 otherwise | $0,\ \forall\tau_1,\tau_2$ |
| $N_{C,V_j}(0,\tau_1,\tau_2)$ | vg $mL^{-1}$ | $0,\ \forall j,\tau_1,\tau_2$ | $0,\ \forall j,\tau_1,\tau_2$ | $0,\ \forall j,\tau_1,\tau_2$ | $0,\ \forall j,\tau_1,\tau_2$ |
| $W(0)$ | cell $mL^{-1}$ | $0,\ \forall j,\tau_1,\tau_2$ | $0,\ \forall j,\tau_1,\tau_2$ | $0,\ \forall j,\tau_1,\tau_2$ | Exp. measurement[S8] |
| $S(0)$ | nmol $mL^{-1}$ | $15\times10^3$ | $15\times10^3$ | $15\times10^3$ | – |
| $T(0)$ | cell $mL^{-1}$ | $1.5\times10^6$ | $1.5\times10^6$ | $1.5\times10^6$ | Exp. measurement[S8] |
| $V_j(0)$ | virus $mL^{-1}$ | $V_1(0)/T(0)=\{10^{-3},10^{-1},1\}$ ;<br>$V_2(0)=0$ | | $0,\ \forall j$ | $V_1(0)/T(0)=\{1,3,30\}$;<br>$V_2(0)/T(0)=\{0,1,3,30\}$ |
| $\Delta\tau$ | hpi | 0.1 | 0.1 | 0.2 | 0.25 |
| Sampling interval | h | 1 | 1 | 1 | 0.5 |
| Model | – | 1 virus | 1 virus | 2 viruses (STV+STV) | (STV+DIP) |
| Parameters | – | Table S3 | Table S3 | Table S4 | Table S5 |

#### Model limitations

The following limitations should be considered when using the simulation framework. The dynamics of substrate consumption and limitation to growth are implemented for only one (generic) nutrient, which is the nutrient that is present at a limiting level in the system. The model can be extended to consider multiple nutrients and metabolites. Further, a distribution with respect to the infection age might exist for the cell growth rate and specific substrate consumption, which are here considered constant constant. However, any model parameter can be converted into functions of the infection age and of the intracellular viral genome level. Although the implementation of such relations in the model would be straightforward, caution must be posed to avoid overfitting and identifiability issues. Case study 4 demonstrates that the current implementation of the model can already explain experimental data with a very satisfactory performance. On the contrary, more parameters might be needed to simulate coinfections from two very different viruses that result in strongly nonlinear coinfection kinetics. The intrinsic model limitation to maximum two viral species can be tackled by lumping together viruses that present similar kinetics. In this scenario, it can be helpful to conduct more simulations, varying the combinations of the multiple viruses that are lumped together into a single viral species. These considerations apply also when more than one type of DIP is present in the system. Unfortunately, the explicit implementation of additional viral species and of their infection age as an additional independent variable would make the model too computationally expensive. Finally, the possibility that infected cells can become uninfected due to cell growth and lysosomal degradation is not explicitly considered. However, the model does consider that these phenomena can reduce the intracellular level of viral genome, even significantly. In these regards, the viral replication rate should be imposed equal to the cell growth rate for viruses that stably integrate into the host genome. On the numerics side, a robust performance and satisfying computational times are achieved in all case studies. This result is not globally guaranteed for very different values of the model parameters. For instance, the default Runge Kutta scheme used for integration of the model equations might not perform as well for systems in which the model parameters induce significant stiffness into the equations. The use of different integration schemes and of automatic differentiation tools would be useful in this situation to improve the simulation performance. Further, adjustments might be made to the convergence tolerances and scaling settings based on the system dynamics and on the mesh size.

### Additional information on Case study 4

Table S7 compares the RSS and the AIC achieved by the model introduced in this work and by the model by Rudiger et al.[S8] in the considered experimental dataset. The AIC is used for model selection as this metric was also used by Rudiger et al.[S8] The RSS is calculated on measurements of (i) STV and (ii) DIP viral titer (Fig. 9), (iii) STV and (iv) DIP intracellular viral genome copy number (Fig. S5), and (v) VCD (Fig. S6). For each variable, residuals are normalized with respect to the maximum measurement in a given experiment, as in Rudiger et al.[S8] Measurements at $t = 0$ are excluded from the RSS computation, as they represent the initial conditions. In experiment 4, no significant dynamics are registered after 40 hours post inoculation. Hence, data collected after 40 hours post inoculation in experiment 4 is not considered for parameter estimation and in the RSS computation, to reduce the simulation time. The AIC is calculated as detailed by Akaike.[S9] The number of estimated parameters used in the AIC computation is 11 for the model presented in this work, and 19 for the model by Rudiger et al.[S8] The model presented in this work is selected as a better model to represent the experimental data, since it achieves lower cumulative RSS and AIC, and lower AIC in 8/12 experiments. Three main features allow our model to outperform the model by Rudiger et al.[S8] First of all, the model by Rudiger et al.[S8] can consider infection of a cell by maximum one unit of the same type of virus (STV or DIP). This is a strong limitation, since the STV MOI and/or the DIP MOI can be at inoculation (or become during the process) larger than 1 (Fig. 9). The model presented in this work can account for the cellular uptake of multiple viruses, without any arbitrary limitation. In our model, the number of viral genomes that reach the nucleus through infection directly affects the viral genome level at the end of replication (Eqs. 10, S19–S22). In turn, the ratio between STV and DIP viral genomes in a coinfected cell affects the STV/DIP ratio in the released progeny (Eq. S14). Further, the level of viral genome affects the death rate of infected cells (Eqs. 4, S5, S7). The temporal difference between subsequent STV and DIP infections is also accurately accounted for in these computations through the calculation of a two-dimensional distribution (Fig. 2). Hence, our model can infer how the infection from more than one STV and/or DIP maps onto the death rate and onto the production of STV/DIP progeny for the infected cells. Instead, the model by Rudiger et al[S8] cannot directly describe the effect of the infection of a cell from more than one STV and/or more than one DIP. Rudiger et al.[S8] improved the fit of their model by introducing 4 extra parameters which affect the rate of transcription, progeny release, and cell growth, based on the STV MOI/DIP MOI ratio at the time of infection of a cell. These heuristic parameters were fit to the experimental data. Our model, instead, directly describes the effect of the STV MOI/DIP MOI ratio by accounting for the uptake of multiple STVs and/or DIPs. Further, the model presented in this work calculates the distribution of infected and coinfected cells and of their intracellular species with respect to the infection ages of both STV and DIP (Fig. 2). Although the infection age from a DIP does not strongly affect the cell cycle nor the viral infection kinetics, a gradient with respect to the DIP infection age can exist for the DIP genome copy number in DIP-infected cells, due to the possibility of re-infection. Gradients of DIP genome copy number in DIP-infected cells can propagate into gradients of STV and DIP genomes copy number with respect to the DIP infection age in coinfected cells, too. Our model considers these phenomena, since, for the reasons mentioned in the previous paragraph, improving the estimation of the viral genomes distribution in the cell population allows to better simulate the STV/DIP competition in coinfected cells. The downside of introducing two-dimensional distributions in our model is the much increased computational burden. However, the choice of lumping together several intermediates into only the most important intracellular species (surface-attached viruses and nuclear viral genomes) and the novel efficient numerical methodology presented here allow our simulation framework to maintain low computational times (Table S1). The novel numeric approach introduced in this work is the third element that allows to outperform the model by Rudiger et al.,[S8] since it allows to compute the infection age distribution with much higher accuracy than the finite difference approach used by Rudiger et al.[S8] (Figs. S1,S2).

Table S7: Case study 4: RSS and AIC achieved by the model introduced in this work and by the model by Rudiger et al.[S8]

| Experiment | RSS | AIC | RSS | AIC |
|---|---|---|---|---|
| | *This work* | | *Rudiger et al.*[S8] | |
| 1 | 4.086 | −31.889 | 2.553 | −29.058 |
| 2 | 8.514 | −45.032 | 6.523 | −40.221 |
| 3 | 12.270 | −94.440 | 31.885 | −13.501 |
| 4 | 10.866 | −56.855 | 24.690 | 1.004 |
| 5 | 3.445 | −24.589 | 1.723 | −25.210 |
| 6 | 8.053 | −36.960 | 4.844 | −40.274 |
| 7 | 9.114 | −39.653 | 4.033 | −57.081 |
| 8 | 13.339 | −24.037 | 10.350 | −18.439 |
| 9 | 2.482 | −32.458 | 1.336 | −31.325 |
| 10 | 9.037 | −32.579 | 190.780 | 99.314 |
| 11 | 8.026 | −44.868 | 5.653 | −43.239 |
| 12 | 7.494 | −47.678 | 4.253 | −54.905 |
| Cumulative | **96.725** | **−739.121** | **288.624** | **−201.642** |

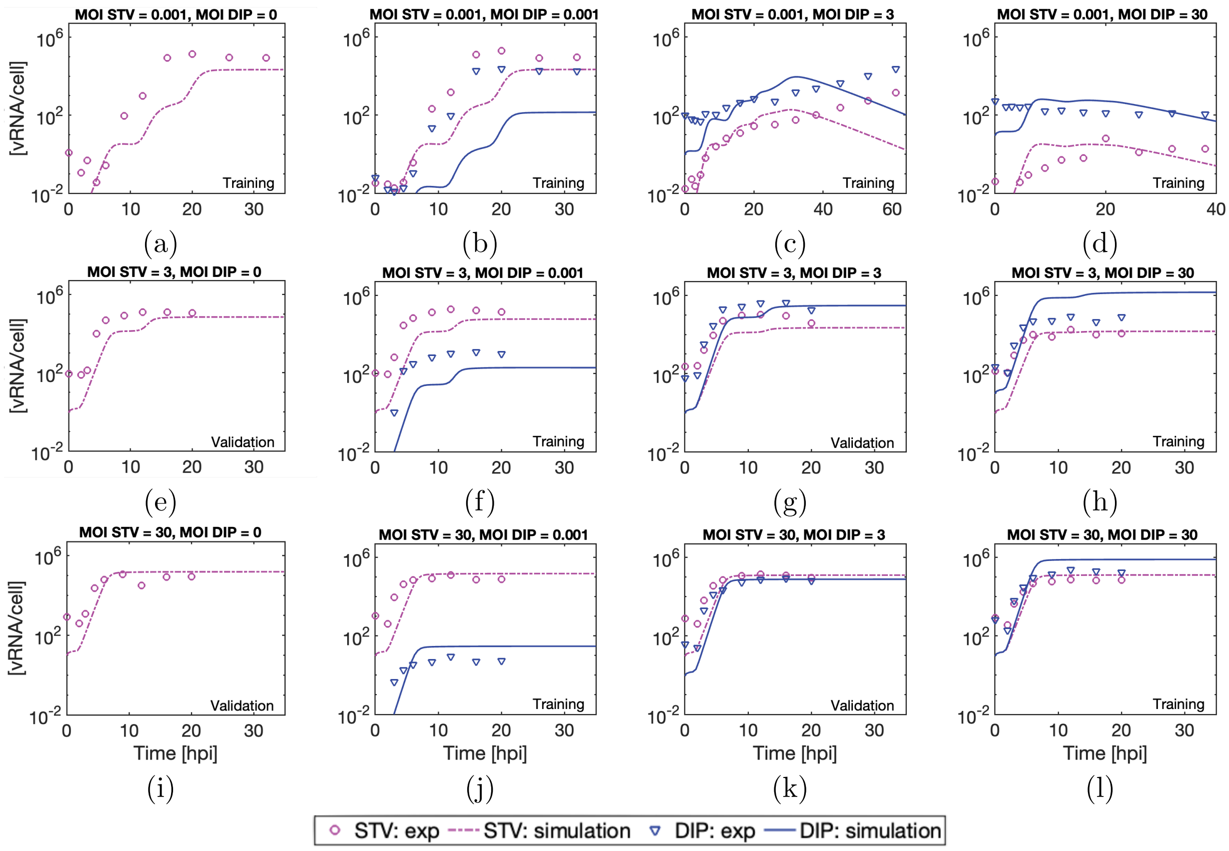


Figure S5: Case study 4: model predictions vs. experimental measurements for the intracellular viral genome concentration in 12 batches of influenza A STV and DIP coinfection. Each batch is inoculated at the reported STV MOI, measured through infectivity assay, and DIP MOI, measured through infectivity assays on MDCK cells genetically modified to express PB2 ("active DIP titer" approach[S10]). Experimental data from Rudiger et al.[S8]

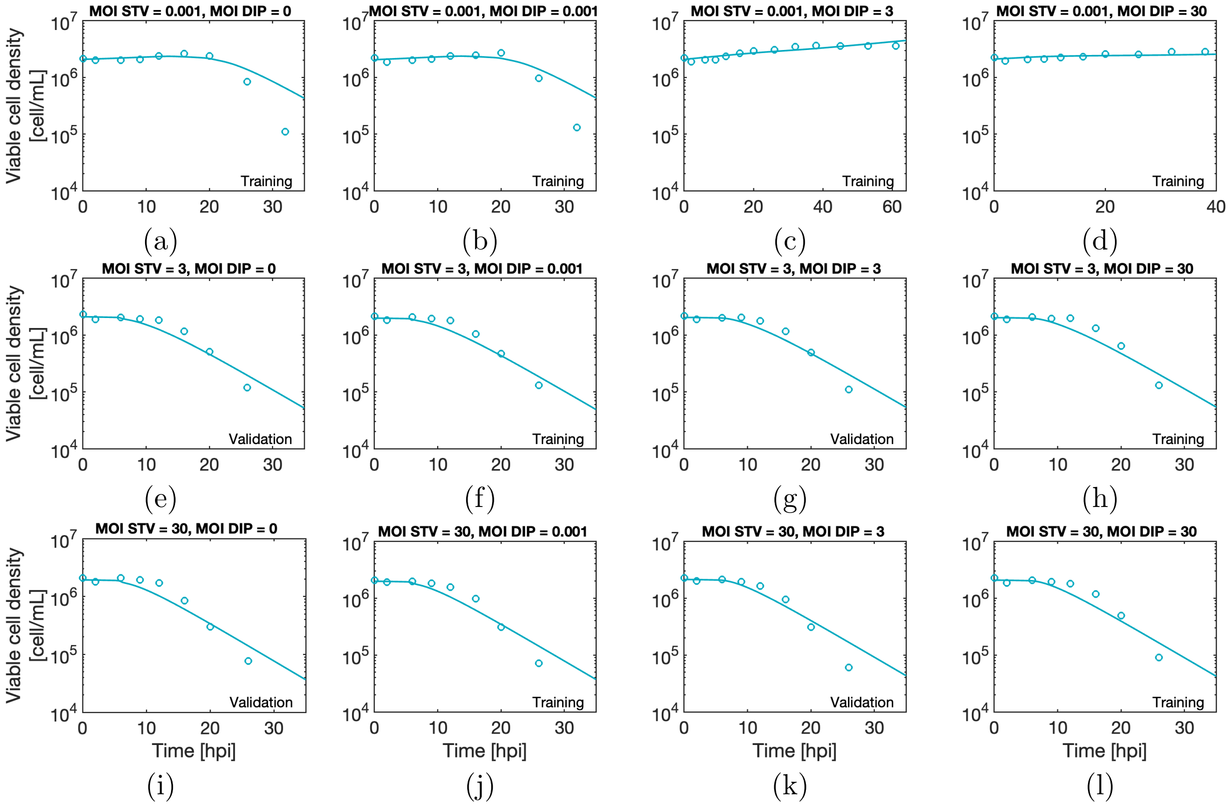


Figure S6: Case study 4: model predictions vs. experimental measurements for the viable cell density of influenza A STV and DIP in 12 batches of influenza A STV and DIP coinfection. Each batch is inoculated at the reported STV MOI, measured through infectivity assay, and DIP MOI, measured through infectivity assays on MDCK cells genetically modified to express PB2 ("active DIP titer" approach[S10]). Experimental data from Rudiger et al.[S8]

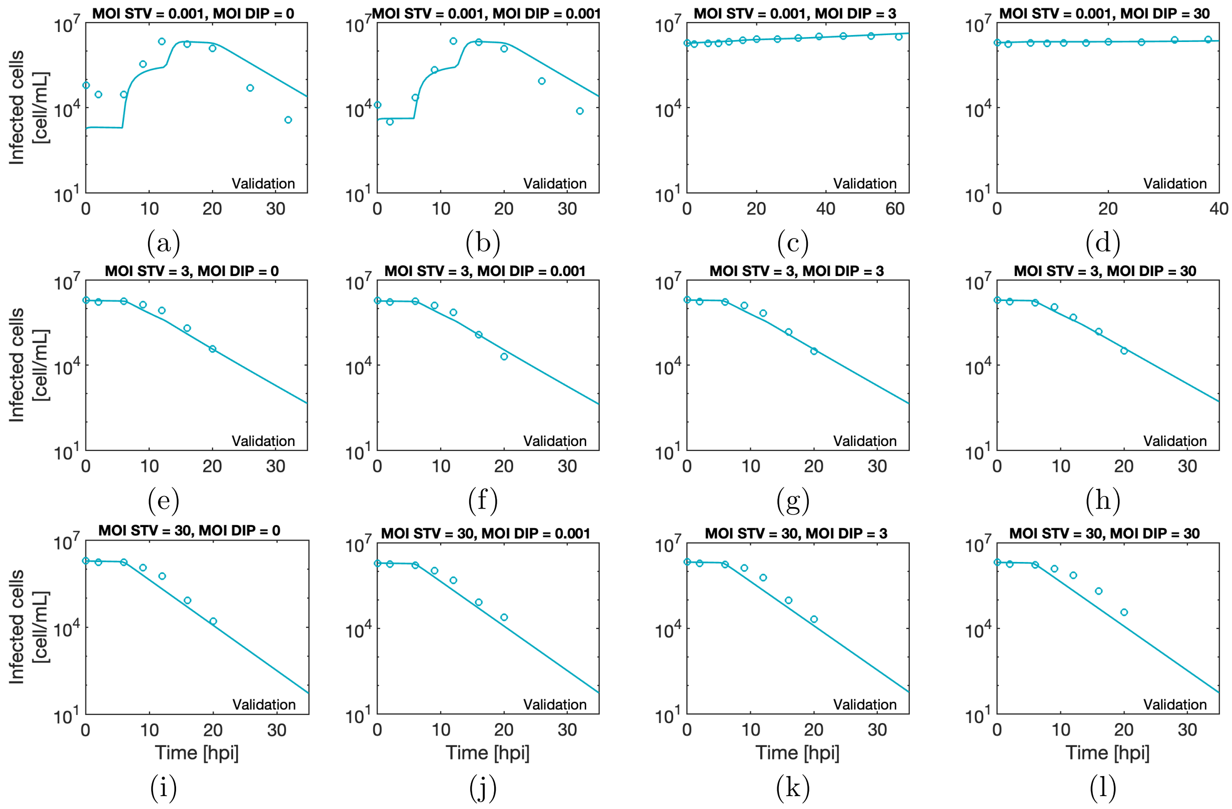


Figure S7: Case study 4: model predictions vs. experimental measurements for the infected cell concentration in 12 batches of influenza A STV and DIP coinfection. The infected cell concentration is the cumulative concentration of cells infective by only one or both STV and DIP. Each batch is inoculated at the reported STV MOI, measured through infectivity assay, and DIP MOI, measured through infectivity assays on MDCK cells genetically modified to express PB2 ("active DIP titer" approach[S10]). Experimental data from Rudiger et al.[S8]

### Supplementary Methods

#### Mathematical model for systems with two viral species

This section presents a novel model for transduction and propagation of two viral species in a suspension culture (Figs. 1–3). The main modeling assumptions are the same posed for systems with only one viral species (Methods section of the main manuscript). The system states and the model parameters are summarized in, respectively, Tables 1 and S2. The model is defined by the set of differential Eqs. S1–S3, S6, S8–S11, S15–S22. The extracellular compartment of the model affects the intracellular compartment by determining the viral uptake dynamics. In turn, the intracellular compartment affects the extracellular level through its influence on both the cell death rate and progeny release, as further described. In this section, all distributed states are introduced as continuous distributions. Throughout the manuscript, continuous distributions are consistently represented using lowercase letters for symbols, while discrete distributions are denoted by the corresponding uppercase letters of the same symbol, with unchanged subscripts and superscripts. Non-distributed states are always reported as uppercase letters. The balance for the uninfected cells concentration $T$

[cell/mL] is:

$$\frac{dT}{dt} = \mu_T T \frac{S}{K_S + S} - T(k_{b,T,V_1} V_1 + k_{b,T,V_2} V_2) - k_{d,T} T + D(T_{\text{in}} - rT), \tag{S1}$$

where $\mu_T$ is the growth kinetic constant for uninfected cells, $S$ [nmol/mL] is the substrate concentration in the system, $K_S$ is the Michaelis-Menten constant for substrate limitation to cell growth, $k_{b,T,V_j}$ is the kinetic constant for binding of virus $j = \{1, 2\}$ to uninfected cells, $k_{d,T}$ is the death kinetic constant for uninfected cells, $D$ is the dilution rate of the system, $T_{\text{in}}$ is the uninfected cells concentration in the feed, and $r$ is the bleeding ratio. Figure 1 clarifies the physical meaning of $D$ and $r$. The terms in the right-hand side of Eq. S1 account for, from left to right: cell growth, infection from $V_1$, infection from $V_2$, cell death, and system inlet/outlet. The balances of $i_k(t, \tau_k)$ [cell mL$^{-1}$ hpi$^{-1}$], the continuous distributions with respect to the infection age $\tau_k$ of the concentrations of cells infected only by virus $k = \{1, 2\}$ are:

$$\frac{\partial i_1}{\partial t} + \frac{\partial i_1}{\partial \tau_1} = \mu_{I_1} i_1 \frac{S}{K_S + S} + k_{b,T,V_1} T V_1 \delta(\tau_1) - i_1 \left( \bar{k}_{d,I_1} + rD + \bar{k}_{b,I_1,V_2} V_2 \right), \tag{S2}$$

$$\frac{\partial i_2}{\partial t} + \frac{\partial i_2}{\partial \tau_2} = \mu_{I_2} i_2 \frac{S}{K_S + S} + k_{b,T,V_2} T V_2 \delta(\tau_2) - i_2 \left( \bar{k}_{d,I_2} + rD + \bar{k}_{b,I_2,V_1} V_1 \right), \tag{S3}$$

where, for $k = \{1, 2\}$, $\tau_k$ is the infection age for virus $k$, $\mu_{I_k}$ is the growth kinetic constant of $i_k$, $\bar{k}_{d,I_k}$ is the death equivalent kinetic constant for $i_k$, $\delta(\cdot)$ is the Dirac delta function, $\bar{k}_{b,I_2,V_1}$ is the binding equivalent kinetic constant of $V_1$ to $i_2$, and $\bar{k}_{b,I_1,V_2}$ is the binding equivalent kinetic constant of $V_2$ to $i_1$. The terms in the right-hand side of Eqs. S2–S3 account for, from left to right: cell growth, infection of uninfected cells, cell death, outlet from the system, and coinfection. The equivalent kinetic constants for binding of virus $j = \{1, 2\}$ to infected cells $i_k$ depend on $\tau_k$, and are calculated, for $k = \{1, 2\}$, as

$$\bar{k}_{b,I_k,V_j}(\tau_k) = \begin{cases} k_{b,T,V_j}, & \text{for } \tau_k < \tau_{b,I_k}, \\ k_{b,T,V_j} \exp(-\beta_{b,I_k}(\tau_k - \tau_{b,I_k})), & \text{for } \tau_k \geq \tau_{b,I_k}, \end{cases} \tag{S4}$$

where $\beta_{b,I_k}$ and $\tau_{b,I_k}$ are suitable coefficients that describe the viral binding downregulation with the progress of the infection age, which is experienced for most viral infections.[S11,S12] Equation S4 assumes that cells infected (only) by virus $k$ undergo binding downregulation for both viruses with the same kinetics, defined by $\beta_{b,I_k}$ and $\tau_{b,I_k}$. However, cells infected by a virus might be prevented from becoming re-infected by the same virus, but not from other (very different) viruses. Equation S4 needs to be updated for simulating systems presenting these characteristics. The death equivalent kinetic parameter depends on both the infection age and the viral genome copy number, and, for $k = \{1, 2\}$, is computed as[S1]

$$\bar{k}_{d,I_k}(t, \tau_k) = \begin{cases} k_{d,T}, & \text{for } \tau_k < \tau_{d,I_k}, \\ k_{d,I_k} \ln(n_{I_k}/i_k) & \text{for } \tau_k \geq \tau_{d,I_k}, \end{cases} \tag{S5}$$

where $k_{d,I_k}$ is the death kinetic constant for cells $i_k$, $\tau_{d,I_k}$ is the infection age at which an increase of death rate is registered for $i_k$, and $n_{I_k}/i_k$ is the viral genome copy number in the cell nucleus, as further discussed. The balance of the coinfected cells concentration $c(t, \tau_1, \tau_2)$ [cell mL$^{-1}$ hpi$^{-1}$], distributed with respect to both infection ages, is

$$\frac{\partial c}{\partial t} + \frac{\partial c}{\partial \tau_1} + \frac{\partial c}{\partial \tau_2} = \mu_C c \frac{S}{K_S + S} + \bar{k}_{b,I_1,V_2} i_1 V_2 + \bar{k}_{b,I_2,V_1} i_2 V_1 - c\left(\bar{k}_{d,C} + rD\right), \tag{S6}$$

where $\mu_C$ is the growth kinetic constant for coinfected cells, and the death equivalent kinetic parameter $\bar{k}_{d,C}$ is calculated as

$$\bar{k}_{d,C}(t, \tau_1, \tau_2) = \begin{cases} k_{d,T}, & \text{for } \tau < \tau_{d,C}, \\ k_{d,C} \ln\left(\frac{\sum_k f_{d,C,V_k} N_{C,V_k}}{c}\right) & \text{for } \tau \geq \tau_{d,C}, \end{cases}, \tag{S7}$$

where $k_{d,C}$ is the death kinetic constant for coinfected cells, $\tau_{d,C}$ is the infection age at which an increase of death rate is registered for coinfected cells, and parameters $f_{d,C,V_1}$ and $f_{d,C,V_2}$ are weighting factors that assume values between 0 and 1, while the argument of the logarithmic function is a weighted sum of the viral genome copy number of virus 1 and virus 2 in the nucleus of coinfected cells. The infection age for coinfected cells is $\tau = \max\{\tau_1, \tau_2\}$ if virus 1 and virus 2 are STVs (Case study 3), while $\tau$ is equal to the infection age from the STV in STV/DIP coinfected cells. For simplicity, the presence of only one substrate $S$ [nmol mL−1] is considered in the default implementation of the model, whose balance is

$$\frac{dS}{dt} = D(S_{\text{in}} - S) - \frac{S}{\phi_S + S}\left(Y_{S,T} T + \int_0^\infty Y_{S,I_1} i_1 \, d\tau_1 + \int_0^\infty Y_{S,I_2} i_2 \, d\tau_2 + \int_0^\infty \int_0^\infty Y_{S,C} c \, d\tau_1 d\tau_2\right), \tag{S8}$$

where $S_{\text{in}}$ is the substrate concentration in the feed, $Y_{S,T}$ is the specific substrate consumption for $T$, $Y_{S,I_1}$ is the specific substrate consumption for $i_1$, $Y_{S,I_2}$ is the specific substrate consumption for $i_2$, $Y_{S,C}$ is the specific substrate consumption for $c$, and the parameter $\phi_S$ describes the kinetics of substrate consumption at low substrate levels, which is fixed to the value of $\phi_S = 0.01$ nmol mL$^{-1}$. A balance is also developed for tracking the concentration of nonviable cells $W$ [cell mL$^{-1}$] in the system:

$$\frac{dW}{dt} = k_{d,T} T + \int_0^\infty \bar{k}_{d,I_1} i_1 \, d\tau_1 + \int_0^\infty \bar{k}_{d,I_2} i_2 \, d\tau_2 + \int_0^\infty \int_0^\infty \bar{k}_{d,c} c \, d\tau_1 d\tau_2 - W(k_{lys} + rD), \tag{S9}$$

where $k_{lys}$ is the lysis kinetic constant for nonviable cells. The balances for the extracellular virion concentration [PFU/mL] for virus 1 ($V_1$) and virus 2 ($V_2$) are, respectively,

$$\begin{aligned}\frac{dV_1}{dt} = &\int_0^\infty (\bar{k}_{v,I_1} - V_1\bar{k}_{b,I_1,V_1}) i_1\, d\tau_1 + \int_0^\infty (\bar{k}_{v,I_2}\phi_{I_2,V_1} - V_1\bar{k}_{b,I_2,V_1}) i_2\, d\tau_2 + \\ &\int_0^\infty\int_0^\infty (\bar{k}_{v,C,V_1} - V_1\bar{k}_{b,C,V_1}) c\, d\tau_1 d\tau_2 - V_1\Big(k_{b,T,V_1}T + k_{d,V_1} + D\Big),\end{aligned} \tag{S10}$$

$$\begin{aligned}\frac{dV_2}{dt} = &\int_0^\infty (\bar{k}_{v,I_1}\phi_{I_1,V_2} - V_2\bar{k}_{b,I_1,V_1}) i_1\, d\tau_1 + \int_0^\infty (\bar{k}_{v,I_2} - V_2\bar{k}_{b,I_2,V_1}) i_2\, d\tau_2 + \\ &\int_0^\infty\int_0^\infty (\bar{k}_{v,C,V_2} - V_2\bar{k}_{b,C,V_1}) c\, d\tau_1 d\tau_2 - V_2\Big(k_{b,T,V_2}T + k_{d,V_2} + D\Big),\end{aligned} \tag{S11}$$

where, for $j = \{1, 2\}$, $\bar{k}_{v,I_j}$ is the progeny release equivalent parameter for cells infected (only) by virus $j$, $\bar{k}_{v,C,V_j}$ is the progeny release equivalent parameter for virus $j$ from coinfected cells, and $\bar{k}_{b,C,V_j}$ is the equivalent kinetic parameter for binding of virus $j$ to coinfected cells. The parameter $\phi_{I_1,V_2}$ ($\phi_{I_2,V_1}$) denotes random generation of $V_2$ ($V_1$) from cells infected only by virus 1 (2); $\phi_{I_1,V_2}$ and $\phi_{I_2,V_1}$ are primarily useful to account for DIP generation from random deletion events. The binding equivalent parameter for coinfected cells is, for $j = \{1, 2\}$,

$$\bar{k}_{b,C,V_j} = \begin{cases} k_{b,T,V_j}, & \text{for } \tau < \tau_{b,C,V_j}, \\ k_{b,T,V_j}\exp(-\beta_{b,C,V_j}(\tau - \tau_{b,C,V_j})), & \text{for } \tau \geq \tau_{b,C,V_j}, \end{cases} \tag{S12}$$

Equivalent parameters $\bar{k}_{v,I_j}$ and $\bar{k}_{v,C,V_j}$ depend on the cell infection age and are respectively calculated as, for $j = \{1, 2\}$:

$$\bar{k}_{v,I_j}(\tau_j) = \begin{cases} 0, & \text{for } \tau_j < \tau^{\text{on}}_{v,I_j} \vee \tau_j > \tau^{\text{off}}_{v,I_j}, \\ k_{v,I_j}, & \text{for } \tau^{\text{on}}_{v,I_j} \leq \tau_j \leq \tau^{\text{off}}_{v,I_j}, \end{cases} \tag{S13}$$

$$\bar{k}_{v,C,V_j}(\tau_1, \tau_2) = \begin{cases} 0, & \text{for } \tau < \tau^{\text{on}}_{v,C} \vee \tau > \tau^{\text{off}}_{v,C}, \\ k_{v,C}\frac{n_{C,V_j}}{n_{C,V_2}+n_{C,V_2}}, & \text{for } \tau^{\text{on}}_{v,C} \leq \tau \leq \tau^{\text{off}}_{v,C} \end{cases} \tag{S14}$$

where $k_{v,I_1}$, $k_{v,I_2}$, and $k_{v,C}$ are the progeny release kinetic constants for, respectively, cells infected only by virus 1, only by virus 2, and for coinfected cells. Equations S13 and S14 approximate the progeny production rate assuming that infected cells release a fixed amount of progeny virus ($k_{v,I_1}$, $k_{v,I_2}$, and $k_{v,C}$; [PFU cell$^{-1}$ h$^{-1}$]) during the infection time interval $[\tau^{\text{on}}_{v,I_j}, \tau^{\text{off}}_{v,I_j}]$ for cells infected by only one virus, and $[\tau^{\text{on}}_{v,C}, \tau^{\text{off}}_{v,C}]$ for coinfected cells. Further, Eq. S14 postulates that coinfected cells release progeny of virus $j = \{1, 2\}$ proportionally to the fraction of viral genome $j$ in the cell nucleus, calculated over the total viral genome copy number for both viruses. Hence, Eq. S14 considers that cells will predominantly release progeny of one viral species if it is prevalent over the other in the cell nucleus (either because of larger number of internalized virions and/or stronger viral amplification). The assumption that virus 1 and 2 compete for releasing their own progeny might not hold true for all combinations of viral species. The parameters of the model can be modified to simulate non-competitive progeny release or other dynamics of interest.

Overall, 8 intracellular species are considered by the model (Fig. 3). For each virus $j = \{1, 2\}$, the model calculates the concentration of virus bound to the surface of infected ($b_{I_j}$) and coinfected ($b_{C,V_j}$) cells, as well as the viral genome in the nucleus of infected ($n_{I_j}$) and coinfected ($n_{C,V_j}$) cells. The concentrations of the intracellular species are computed with respect to the whole volume of the system [# mL$^{-1}$ hpi$^{-1}$], and they are distributed along the infection age(s). Under the deterministic modeling assumptions followed here, uninfected cells that become infected by virus 1 at the same time instant will inherently present the same time profiles of $\tau_1$, $b_{I_1}$, and $n_{I_1}$, until cell death or coinfection occur. The same applies to uninfected cells that become infected by (only) virus 2 at the same time instant. Further, viable coinfected cells that have both the same $\tau_1$ and $\tau_2$ will also present the same time profiles of $b_{C,V_1}$, $b_{C,V_2}$, $n_{C,V_1}$, and $n_{C,V_2}$. Hence, the distributions of intracellular species concentration with respect to the infection age(s) directly map to the distribution of the infected and coinfected cell populations with respect to the infection age. It directly follows that to convert the intracellular species concentration in a per cell basis [#/cell], $b_{I_1}(t, \tau_1)$ and $n_{I_1}(t, \tau_1)$ should be normalized by $i_1(t, \tau_1)$, while $b_{I_2}(t, \tau_2)$ and $n_{I_2}(t, \tau_2)$ should be normalized by $i_2(t, \tau_2)$. For coinfected cells, instead, $b_{C,V_1}(t, \tau_1, \tau_2)$,$b_{C,V_2}(t, \tau_1, \tau_2)$,$n_{C,V_1}(t, \tau_1, \tau_2)$ and $n_{C,V_2}(t, \tau_1, \tau_2)$ should all be normalized by $c(t, \tau_1, \tau 2)$. The mass balances for the intracellular species are

$$\frac{\partial b_{I_1}}{\partial t} + \frac{\partial b_{I_1}}{\partial \tau_1} = V_1\left(k_{b,T,V_1}T\delta(\tau_1) + \bar{k}_{b,T,V_1}i_1\right) - b_{I_1}\left(\bar{k}_{d,I_1} + rD + \bar{k}_{b,I_1,V_2}V_2 + k_{i,I_1}\right), \quad \text{(S15)}$$

$$\frac{\partial b_{I_2}}{\partial t} + \frac{\partial b_{I_2}}{\partial \tau_2} = V_2\left(k_{b,T,V_2}T\delta(\tau_2) + \bar{k}_{b,T,V_2}i_2\right) - b_{I_2}\left(\bar{k}_{d,I_2} + rD + \bar{k}_{b,I_2,V_1}V_1 + k_{i,I_2}\right), \quad \text{(S16)}$$

$$\begin{aligned}\frac{\partial b_{C,V_1}}{\partial t} + \frac{\partial b_{C,V_1}}{\partial \tau_1} + \frac{\partial b_{C,V_1}}{\partial \tau_2} = {} & \bar{k}_{b,I_1,V_2}V_2 b_{I_1}\delta(\tau_2) + \bar{k}_{b,I_2,V_1}V_1 i_2\delta(\tau_1) + \bar{k}_{b,C,V_1}cV_1 - \\ & b_{C,V1}\left(\bar{k}_{d,C} + rD + \bar{k}_{i,C,V_1}\right),\end{aligned} \quad \text{(S17)}$$

$$\begin{aligned}\frac{\partial b_{C,V_2}}{\partial t} + \frac{\partial b_{C,V_2}}{\partial \tau_1} + \frac{\partial b_{C,V_2}}{\partial \tau_2} = {} & \bar{k}_{b,I_2,V_1}V_1 b_{I_2}\delta(\tau_1) + \bar{k}_{b,I_1,V_2}V_2 i_1\delta(\tau_2) + \bar{k}_{b,C,V_2}cV_2 - \\ & b_{C,V_2}\left(\bar{k}_{d,C} + rD + \bar{k}_{i,C,V_2}\right),\end{aligned} \quad \text{(S18)}$$

$$\frac{\partial n_{I_1}}{\partial t} + \frac{\partial n_{I_1}}{\partial \tau_1} = \eta_{I_1}k_{i,I_1}b_{I_1} + \bar{k}_{r,I_1}n_{I_1} - n_{I_1}\left(\bar{k}_{d,I_1} + k_{d,N_1} + rD + \bar{k}_{b,I_1,V_2}V_2\right), \quad \text{(S19)}$$

$$\frac{\partial n_{I_2}}{\partial t} + \frac{\partial n_{I_2}}{\partial \tau_2} = \eta_{I_2}k_{i,I_2}b_{I_2} + \bar{k}_{r,I_2}n_{I_2} - n_{I_2}\left(\bar{k}_{d,I_2} + k_{d,N_2} + rD + \bar{k}_{b,I_2,V_1}V_1\right), \quad \text{(S20)}$$

$$\frac{\partial n_{C,V_1}}{\partial t} + \frac{\partial n_{C,V_1}}{\partial \tau_1} + \frac{\partial n_{C,V_1}}{\partial \tau_2} = \eta_{C,V_1} k_{i,C,V_1} b_{C,V_1} + \bar{k}_{r,C,V_1} n_{C,V_1} + \bar{k}_{b,I_1,V_2} n_{I_1} V_2 \delta(\tau_2) - n_{C,V_1}\left(\bar{k}_{d,C} + k_{d,N_1} + rD\right), \tag{S21}$$

$$\frac{\partial n_{C,V_2}}{\partial t} + \frac{\partial n_{C,V_2}}{\partial \tau_1} + \frac{\partial n_{C,V_2}}{\partial \tau_2} = \eta_{C,V_2} k_{i,C,V_2} b_{C,V_2} + \bar{k}_{r,C,V_2} N_{C,V_2} + \bar{k}_{b,I_2,V_1} n_{I_2} V_1 \delta(\tau_1) - n_{C,V_2}\left(\bar{k}_{d,C} + k_{d,N_2} + rD\right). \tag{S22}$$

Equations S15–S22 are developed with the same assumptions used for deriving Eqs. 9 and 10 and present equivalent contributions. Equations S15–S22 also account for an additional contribution that moves the intracellular content of infected cells that become coinfected to the mass balance of the relevant intracellular species in coinfected cells. The replication equivalent kinetic constants for infected and coinfected cells are calculated as

$$\bar{k}_{r,I_k}(\tau_k) = \begin{cases} 0, & \text{for } \tau_k < \tau_{r,I_k}^{\text{on}} \vee \tau_k > \tau_{r,I_k}^{\text{off}}, \\ k_{r,I_k}, & \text{for } \tau_{r,I_k}^{\text{on}} \leq \tau_k \leq \tau_{v,I_k}^{\text{off}}. \end{cases} \tag{S23}$$

$$\bar{k}_{r,C,V_j}(\tau_1, \tau_2) = \begin{cases} 0, & \text{for } \tau < \tau_{r,C}^{\text{on}} \vee \tau > \tau_{r,C}^{\text{off}}, \\ k_{r,C,V_j}, & \text{for } \tau_{r,C}^{\text{on}} \leq \tau \leq \tau_{r,C}^{\text{off}}. \end{cases} \tag{S24}$$

Equation S24 assumes that viral genome amplification always occurs in coinfected cells with the same kinetics, indipendently on whether a cell is predominantly infected by one virus or the other. Since the model computes the distribution of viral genomes copy number in coinfected cells, it is straightforward to implement more complicated dynamics in Eq. S24, if of interest for certain applications.

#### Comparison with the solution derived from the method of characteristics

The solution from the method of characteristics reported in Figs. S1–S2 is obtained as follow. The model for systems with only one virus is considered (Eqs. 1, 2, 5, 7–10). The distributed states $i(t,\tau)$, $b(t,\tau)$, and $n(t,\tau)$ are discretized along $M$ characteristic lines of infection age $\tau^m$ into, respectively, $B(t,\tau^m) = B^m$, $I(t,\tau^m) = I^m$, and $N(t,\tau^m) = N^m$, for $m = 1, 2, \ldots, M$. The PDEs of the model (Eqs. 2, 9–10) are converted into the ODEs

$$\frac{dI^m}{dt} = \mu_I I^m \frac{S}{K_S + S} + k_{b,T} T V \delta_{m1} - I^m(\bar{k}_{d,I} - rD), \tag{S25}$$

$$\frac{dB^m}{dt} = V(k_{b,T} T \delta_{m1} + \bar{k}_{b,I} I^m) - B^m(k_i + rD + \bar{k}_{d,I}), \tag{S26}$$

$$\frac{dN^m}{dt} = \eta k_i B^m + N^m(\bar{k}_{d,I} - k_{d,N} + \bar{k}_r + rD), \tag{S27}$$

$$\frac{d\tau^m}{dt} = 1, \tag{S28}$$

for $m = 1, 2, \ldots, M$. The integrals in Eqs. 5, 7, 8 are converted into sums across all characteristic lines. The system of ODEs and PDEs of Eqs. 1,2, 5, 7–10 becomes the system of ODEs of Eqs. 1, 5, 7, 8, S25–S28, which are integrated with a Runge-Kutta scheme with adaptive time stepping. The model is initialized with only one characteristic line ($m = 1$), with infection age $\tau^1 = 0$. At the beginning of each integration step, a new characteristic line with infection age equal to 0 is created. This approach solves the PDEs of the model with high accuracy, since it computes the distributed states with a fine mesh defined by the time series of the integration steps, and the increase of infection age of infected cells is explicitly computed during integration (Eq. S28). However, creating new characteristic lines at every integration step can become prohibitively expensive in continuous processing and, more generally, for all systems with two viruses, due to the large number of combinations of infection ages that would be needed to describe the two-dimensional distributions associated to coinfected cells. Further, numerical instability was observed due to the intrinsic poor scaling of a state vector in which the infected cell concentration is discretized with integration steps that can vary of orders of magnitude.

The integration-reallocation approach introduced in this work (see Methods section of the main manuscript) takes several of the advantages of the method of characteristics, but groups together infected cells (and the respective intracellular species) based on a fixed mesh, rather than creating a new characteristic line at every integration step. The reallocation step is equivalent to solving Eq. S28 at the discrete time steps defined by the mesh spacing $\Delta\tau$. As a result, the numeric methodology here introduced allows to track the infection front accurately (Figs. S1–S2), but at a much lower computational cost than the method of characteristics.[S13]

#### Comparison with the solution derived from finite differences

The solution from the finite differences approach reported in Figs. S1–S2 is obtained through the following upwind scheme. The model for systems with only one virus is considered (Eqs. 1, 2, 5, 7–10). The infection age $\tau$ is discretized into $K$ nodes. All nodes have size $\Delta\tau$, except for node $K$, which corresponds to the interval $[\tau_{\max}, \infty)$ in the original space of the continuous variable $\tau$. Mesh nodes $k = 1, 2, 3, \ldots, L$ have infection age $\tau^k$=0, $\Delta\tau$, $2\Delta\tau$, $\ldots$, $\tau_{\max}$. The distributed states $b(t,\tau)$, $i(t,\tau)$, and $n(t,\tau)$ are discretized with respect to $\tau$ into the sets of $K$ states denoted as, respectively, $B(t,\tau^k) = B^k$, $I(t,\tau^k) = I^k$, and $N(t,\tau^k) = N^k$, for $k = 1, 2, \ldots, K$. The PDEs of the model (Eqs. 2, 9–10) are discretized into the ODEs:

$$\frac{dI^k}{dt} = \mu_I I^k \frac{S}{K_S+S} - \frac{I^k - I^{k-1}}{\Delta\tau} + k_{b,T}TV\delta_{k1} - I^k(\bar{k}_{d,I} - rD), \text{ for } k = 1,2,3,\ldots,K-1, \tag{S29}$$

$$\frac{dI^K}{dt} = \mu_I I^K \frac{S}{K_S+S} + \frac{I^{K-1}}{\Delta\tau} - I^K(\bar{k}_{d,I} - rD), \tag{S30}$$

$$\frac{dB^k}{dt} = V(k_{b,T}T\delta_{k1} + \bar{k}_{b,I}I^k) - \frac{B^k - B^{k-1}}{\Delta\tau} - B^k(k_i + rD + \bar{k}_{d,I}), \text{ for } k = 2,3,\ldots,K-1, \tag{S31}$$

$$\frac{dB^K}{dt} = V + \bar{k}_{b,I}I^K \frac{B_{K-1}}{\Delta\tau} + \bar{k}_{b,I}I^K) - B^K(k_i + rD + \bar{k}_{d,I}), \tag{S32}$$

$$\frac{dN^k}{dt} = \eta k_i B^k - \frac{N^k - N^{k-1}}{\Delta\tau} + N^k(\bar{k}_{d,I} - k_{d,N} + \bar{k}_r + rD), \text{ for } k = 2,3,\ldots,K-1, \tag{S33}$$

$$\frac{dN^K}{dt} = \eta k_i B^K + \frac{N^{K-1}}{\Delta\tau} + N^K(\bar{k}_{d,I} - k_{d,N} + \bar{k}_r + rD). \tag{S34}$$

The integrals in Eqs. 5, 7, 8 are converted into sums across all the mesh nodes. The system of ODEs and PDEs of Eqs. 1, 2, 5, 7–10 becomes the system of ODEs of Eqs. 1, 5, 7, 8, S29–S34, which are integrated with a Runge-Kutta scheme with adaptive time stepping.

#### Comparison with the lumped parameter model

The lumped parameter model used for the simulations in Figs. S3 and S4 is defined by the ODEs[S14]

$$\frac{dT}{dt} = \mu_T T \frac{S}{K_S+S} - \hat{k}_b\, TV - k_{d,T}T + D(T_{\text{in}} - rT), \tag{S35}$$

$$\frac{dV}{dt} = \hat{k}_v I - \hat{k}_b V(T+I) - k_{d,V}V - DV, \tag{S36}$$

$$\frac{dI}{dt} = \mu_I I \frac{S}{K_S+S} + \hat{k}_b TV - I(\hat{k}_{d,I}I + rD), \tag{S37}$$

$$\frac{dS}{dt} = D(S_{\text{in}} - S) - \frac{S}{\phi_S + S}(Y_{S,T}T + Y_{S,I}I), \tag{S38}$$

Equations S35–S38 are equivalent to Eqs. 1, 2, 5, and 7 of the distributed model introduced in this work, except that parameters $\hat{k}_b$, $\hat{k}_v$, and $\hat{k}_{d,I}$ are constant in the lumped parameter model, instead of being dependent on the infection age (Eqs. 3, 6) and on the viral genome copy number (Eq. 4). For a fair comparison, the parameters of the lumped parameter model that are physically equivalent to those of the distributed parameter model ($\mu_T$, $K_S$, $k_{d,T}$, $k_{d,V}$, $\mu_I$, $\phi_S$, $Y_{S,T}$, and $Y_{S,I}$) are taken as in the distributed parameter model (Table S3). Parameters $\hat{k}_b$, $\hat{k}_v$, and $\hat{k}_{d,I}$ are, instead, estimated through maximum likelihood estimation from synthetic measurements of $T$, $V$, $I$, and $S$ generated through the distributed parameter model and reported in Fig. S3. In the maximum likelihood estimation routine, measurements of a given variable are normalized by the maximum value that the variable assumes in the dataset. The estimated parameters ($\hat{k}_b = 2.067 \times 10^{-9}$ mL cell$^{-1}$ h$^{-1}$, $\hat{k}_v = 5.49$ PFU cell$^{-1}$

$\text{h}^{-1}$, and $\hat{k}_{d,I} = 0.062\ \text{h}^{-1}$) are used in the simulations reported in Figs. S3 and S4. Despite the re-estimation of $\hat{k}_b$, $\hat{k}_v$, and $\hat{k}_{d,I}$, the lumped parameter model fails to reproduce the characteristic dynamics of baculovirus infection.
